# “Two-Birds-One-Stone” Oral Nanotherapeutic Designed to Target Intestinal Integrins and Regulate Redox Homeostasis for Ulcerative Colitis Treatment

**DOI:** 10.1101/2024.01.02.573980

**Authors:** Long Huang, Wei Hu, Long Qun Huang, Qin Xuan Zhou, Zheng Yang Song, Heng Yu Tao, Bing Xu, Can Yang Zhang, Yi Wang, Xin-Hui Xing

## Abstract

Designing highly efficient orally administrated nanotherapeutics with specific inflammatory site-targeting functions in the gastrointestinal (GI) tract for ulcerative colitis (UC) management is a significant challenge. Straightforward and adaptable modular multifunctional nanotherapeutics represent groundbreaking advancements and are crucial to promoting broad application in both academic research and clinical practice. In this study, we focused on exploring a specific targeting modular and functional oral nanotherapy, serving as “one stone”, for the directed localization of inflammation and the regulation of redox homeostasis, thereby achieving effects against “two birds” for UC treatment. The designed nanotherapeutic agent OPNs@LMWH, which has a core-shell structure composed of oxidation-sensitive ε-polylysine nanoparticles (OPNs) in the core and low-molecular-weight heparin (LMWH) in the shell, exhibited specific active targeting effects and therapeutic efficacy simultaneously. We qualitatively and quantificationally confirmed that OPNs@LMWH possessed high integrin αM-mediated immune cellular uptake efficiency and preferentially accumulated in inflamed lesions. Compared with bare OPNs, OPNs@LMWH exhibited enhanced intracellular reactive oxygen species (ROS) scavenging and anti-inflammatory effects. After oral administration of OPNs@LMWH to mice with dextran sulfate sodium (DSS)-induced colitis, robust resilience was observed. OPNs@LMWH effectively ameliorated oxidative stress and inhibited the activation of inflammation-associated signalling pathways while simultaneously bolstering the protective mechanisms of the colonic epithelium. Overall, these findings underscore the compelling dual functionalities of OPNs@LMWH, which enable effective oral delivery to inflamed sites, thereby facilitating precise UC management.

## Introduction

The global incidence and prevalence of ulcerative colitis (UC), a chronic remitting-relapsing inflammatory disorder of the large intestine, have been increasing rapidly worldwide, yet this disease is incurable and has a multifaceted pathophysiology^1–3^. The European Crohn’s and Colitis Organization (ECCO) and American Gastroenterological Association (AGA) recommend that patients follow a step-up approach with chemical drugs or monoclonal antibodies to achieve the desired remission because there is no known clinically available or effective cure for UC^4–6^. However, the frequent and long-term use of conventional small-molecule therapeutics or biologic-based immunosuppressive drugs can lead to severe complications, such as opportunistic infections, malignancies, autoimmunity, and liver toxicity^7–9^. Therefore, designing and developing effective therapeutics and treatment strategies for UC is very important, which may be achieved based on active component discovery and precise delivery by drawing inspiration from the evidence-based understanding of inflammatory signalling and drug delivery methods^10^.

In prior research, precise drug delivery has been identified as a significant challenge^11,12^. The success of precise delivery is fundamentally dependent on the judicious choice of targeting strategy. A variety of strategies have been devised to target intestinal inflammation sites, such as microbial enzyme degradation, pH adjustment, and mucosal charge reversal at inflamed locations^13–17^. However, the complex and ever-changing environment of gastrointestinal diseases significantly undermines the reliability and precision of UC treatment outcomes^18^. Given the propensity of immune cells to accumulate at disease sites, the use of ligand molecules to target immune cells seems more promising. Specific immune cells, such as monocytes, innate lymphoid cells, and T cells, in addition to various associated regulatory cytokines, have been extensively highlighted in the context of inflammatory bowel disease^19–21^. Furthermore, studies have pinpointed the altered trafficking of immune cell circuits as key drivers of mucosal inflammation and tissue destruction in UC^22^, likening it to a severe “traffic jam” at lesion sites. Consequently, integrins such as α4β7 and αMβ2, which have been implicated in immune cell-related adhesion and signalling, could serve as excellent targets^23^. Recently, more studies have tended to focus on interference with integrin function^24^, however, few studies have explored functional integrin-targeting molecules suitable for the complex gastrointestinal environment of UC. As a result, there is scant research on oral therapeutic systems that can specifically target inflammatory cell infiltration via integrins.

When contemplating this issue, we thought of heparin. Heparin is an anionic polysaccharide comprising highly sulfated disaccharide units. Low-molecular-weight heparin (LMWH), prepared by the depolymerization of unfractionated heparin (UFH), has been recognized as an attractive anticoagulant and has been successfully used in clinical settings for several decades^25,26^. Intriguingly, due to the complex sequence of disaccharide units, where anticoagulant fragments account for only 30 % of the polysaccharide^27^, LMWH has been demonstrated to have potential beyond its anticoagulant properties and has been reported to be effective in the treatment of many other diseases, including infectious diseases, lung cancer and Alzheimer’s disease^27–29^. More importantly, heparin reportedly serves as an adhesive ligand for integrins α4 and αM^23,30,31^, known as CD49d and CD11b, respectively. Inspired by these findings, we hypothesize that LMWH via oral administration could be a promising inflammatory site-targeting ligand for developing precise oral therapy for UC.

Since conventional immunosuppressive drugs for UC management fail to fully address treatment needs, it is crucial to develop groundbreaking strategies with multiple therapeutic effects for rebalancing intestinal disorders in UC patients. Reactive oxygen species (ROS), as integral factors affecting intestinal homeostasis, are signalling molecules extensively involved in numerous cellular processes^32^. However, the disproportionate generation of ROS during the UC pathogenesis process, which can lead to an increase by a factor of 10 to 100^33^, inflicts severe tissue damage on the intestinal epithelium and triggers the overproduction of inflammatory mediators. While small-molecule drugs have been found to have unavoidable limitations^34,35^, there is mounting evidence that biocompatible polymer-based materials, typically those based on glucose-related polysaccharides, show significant potential in suppressing oxidative stress and inflammatory mediators, thereby regulating the redox balance in the colon^9,36–40^. Unfortunately, the interactions of multiple hydroxy groups on the glucose residues of polysaccharides introduce structural uncertainty. To design optimal ROS scavenging materials with a defined structure and well-controlled function, alternative biocompatible polymers need to be explored. For instance, ε-polylysine (EPL), a cationic and biodegradable polypeptide produced by microbial fermentation, has garnered increased interest in food chemistry and oral delivery systems^41^. The presence of a single amino group on each repeating unit of EPL enables an available defined structure.

Here, we strongly emphasize the importance of creating precise nanotherapies that integrate multiple functions. Therefore, we propose a modular orally administrated nanotherapeutic consisting of oxidation-sensitive ε-polylysine (OsEPL) nanoparticles (OPNs) as the core, positively charged chitosan (CS) as the middle layer and negatively charged LMWH as the outermost layer to achieve “two-birds-one-stone” treatment of UC. Considering the adaptability and prospective engineering applications of these materials, we utilized nanoprecipitation and layer-by-layer self-assembly methodologies in the fabrication process. We hypothesize that orally administrated nanoparticles can not only target integrin-associated immune cell recruitment and trafficking through the specific interaction between LMWH and integrin αM but also regulate intestinal redox homeostasis through the ROS scavenging function of OsEPL. Overall, we expect that our OPNs@LMWH may be translated to topical UC therapy with high patient compliance *via* oral administration with high therapeutic efficacy and few side effects. Moreover, the strategy of “two-birds-one-stone” might be straightforward and effective for dealing with complex clinical problems.

## Results

### Preparing integrin-targeting OPNs@LMWH by nanoprecipitation and a layer-by-layer process

The oxidation-sensitive material (OsEPL) was designed and synthesized by introducing a hydrophobic phenylboronic ester group that is sensitive to ROS into the side chain amino group, as shown in Supplementary Scheme 1 and Supplementary Fig. 1. The structures of the intermediate and OsEPL were confirmed by ^1^H nuclear magnetic resonance (^1^H-NMR) (Supplementary Figs. 2-4), indicating successful synthesis. Additionally, we constructed a series of OsEPLs with different feeding ratios of phenylboronic acid pinacol ester to amino moiety. As the molar ratio of the hydrophobic side chain increased, the OsEPL gradually formed an amphiphilic polymer in the aqueous phase. In addition, with a max feeding ratio, OsEPL underwent rapid disintegration when incubated with peroxide (Supplementary Fig. 5). Then, we obtained OPNs and OPNs@LMWH by nanoprecipitation and layer-by-layer methods, as illustrated in Fig. 1a. To obtain nanoparticles coated with LMWH, the OPNs were dispersed in CS buffer, which helped to form nanoparticles with a positive charge (OPNs@CS), followed by incubation with an aqueous solution of LMWH to prepare OPNs@LMWH. The OPNs@LMWH presented a spherical core-shell structure with an average particle size of approximately 600 nm and a low polydispersity index (PDI; less than 0.2 on average) according to scanning electron microscopy (SEM) (Fig. 1b), transmission electron microscopy (TEM) (Fig. 1b, c) and dynamic light scattering (DLS) analyses (Fig. 1d). We also observed charge reversal between the OPNs, OPNs@CS and OPNs@LMWH (Fig. 1e), which indicated successful coating. Considering the goal of using this nanotherapeutic for oral administration, its stability within the gastrointestinal tract is a matter of concern. The stability of OPNs@LMWH was investigated by incubation with both dd-H_2_O (4 °C) and simulated gastric/intestinal fluids (SGF and SIF) (Supplementary Fig. 7a). Our results indicated that the OPNs@LMWH had excellent stability both during storage (Supplementary Fig. 7b) and after oral administration (Supplementary Fig. 7c-e).

**Fig. 1.**
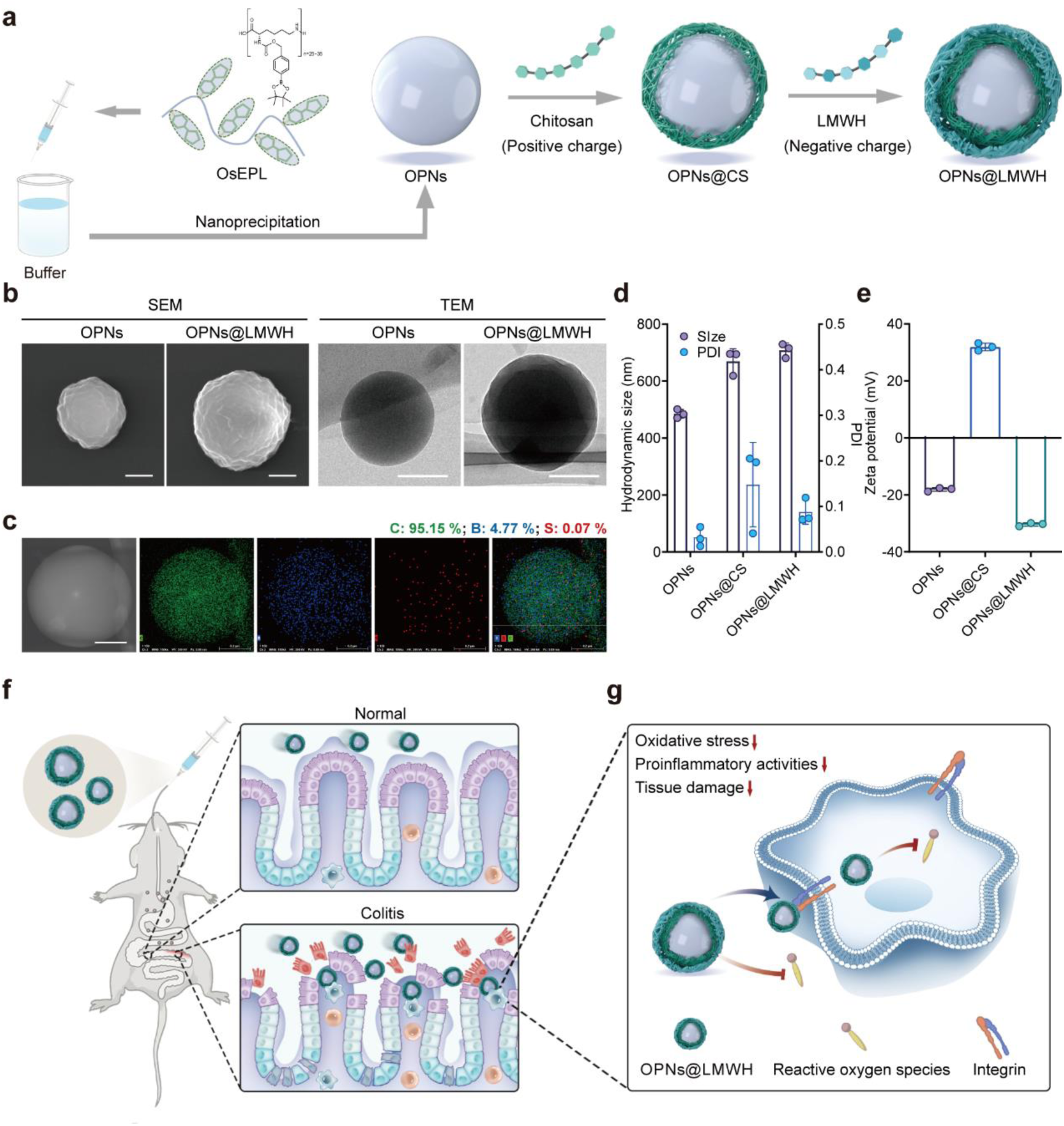
Preparation and characterization of OPNs@LMWH. **a,** Schematic illustration of the preparation of OPNs@LMWH. **b,** Representative SEM and TEM images of OPNs and OPNs@LMWH. Scale bar, 200 nm. **c,** The elements C, B, and S in OPNs@LMWH. Scale bar, 200 nm. **d, e,** Particle size (**d**) and zeta potential (**e**) measurements of different nanoparticles determined by DLS. The data are presented as the means ± SDs. **f, g,** Schematic of the oral administration of OPNs@LMWH targeting immune cell recruitment and regulating intestinal redox homeostasis. OPNs@LMWH, when administered orally, can accumulate at the site of colitis inflammation (**f**). OPNs@LMWH efficiently regulate redox homeostasis by targeting the inflammation-related integrin (**g**).

### Internalization by M1 macrophages and accumulation in inflamed colon tissue infiltrated by inflammatory cells

After the preparation procedures were established, we next compared the ability of OPNs@LMWH and OPNs to be taken up by proinflammatory immune cells. To mimic the interactions between nanoparticles and proinflammatory immune cells at the lesion site, RAW264.7 cells were activated by lipopolysaccharides (LPS) and IFN-γ to form M1 macrophages, which were subsequently incubated with fluorescein amine (FA)-labelled OPNs (FA-OPNs) or FA-labelled OPNs@LMWH (FA-OPNs@LMWH). The samples were subsequently imaged via confocal laser scanning microscopy (CLSM), as shown in Fig. 2a and Supplementary Fig. 8. Compared with the OPN sample (untreated), the OPNs@LMWH sample showed a much greater green fluorescence intensity, indicating significantly enhanced cellular uptake efficiency. Then, we pretreated RAW264.7 cells with an antibody against the known receptor integrin αM to interfere with the potential interaction between LMWH and integrin αM. Consistent with our expectations, reduced attachment of OPNs@LMWH was observed (Fig. 2a, b). For further illustration, we confirmed that free fluorescein was not taken up by cells, while both OPNs and OPNs@LMWH were taken up after 8 h of incubation, which suggested that the enhanced cellular uptake efficiency relied on the LMWH coating layer (Supplementary Fig. 9).

**Fig. 2.**
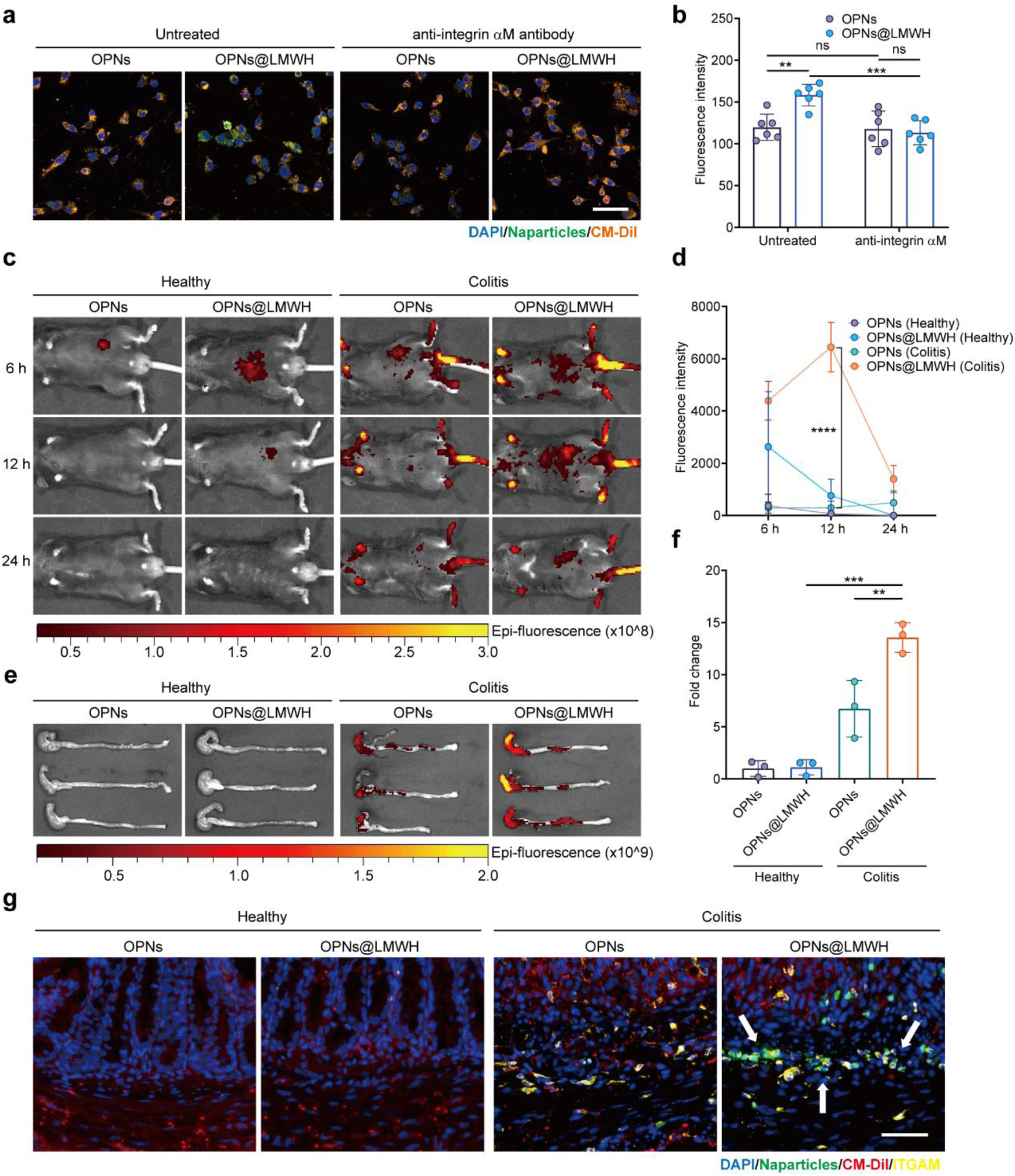
OPNs@LMWH were internalized by M1 macrophages and accumulated in inflamed colon tissue infiltrated by inflammatory cells. **a,** Activated M1-type RAW264.7 cells were incubated with FA-OPNs or FA-OPNs@LMWH after pretreatment with or without the anti-integrin αM antibody, and visualized via CLSM. Scale bar, 50 μm. **b,** Quantitative analysis of the FA fluorescence intensity in images taken by CLSM. **c, d,** Representative healthy and colitis model mice were orally administered ICG-OPNs or ICG-OPNs@LMWH, imaged by an in vivo imaging system (IVIS) and (**c**) the ICG fluorescence signal around the abdomen was quantified (**d**). **e, f,** After 24 h, the colons were obtained and imaged with an IVIS (**e**), after which the ICG fluorescence signal was quantified (**f**). **g,** Healthy or colitis model mice were orally administered FA-OPNs or FA-OPNs@LMWH, and colon tissues were collected after 24 h and stained with DAPI, CM-Dil and integrin αM for visualization via CLSM. Scale bar, 50 μm. Representative images or quantitative analysis of n=6 biologically independent samples (**a, b**) and n=3 animals from two independent experiments (**c-g**) are shown. The data are presented as the means ± SDs. * *p* < 0.05, ** *p* < 0.01, *** *p* < 0.001, **** *p* < 0.0001 and ns (not significant) *p* > 0.05, analysed by two-way ANOVA (**b, d**) or ordinary one-way ANOVA (**f**) for multiple comparisons.

Administering drinking water containing dextran sulfate sodium (DSS) is a well-recognized method to construct a murine model that simulates human UC. We generated DSS-induced colitis model mice to study the in vivo distribution of orally administered OPNs@LMWH. Since we successfully confirmed that OPNs@LMWH targeted integrin αM in vitro, we subsequently confirmed that integrin αM was enriched in colitis model mice because of immune cell recruitment to the colonic submucosal layer (Supplementary Fig. 10). As expected, we successfully observed high accumulation of OPNs@LMWH at the lesion site in vivo. After oral administration of indocyanine green (ICG)-labelled OPNs@LMWH (ICG-OPNs@LMWH), ICG-OPNs@LMWH presented a prolonged retention time in vivo in colitis model mice, and the fluorescence signal peaked at 12 h (Fig. 2c, d). Compared with OPNs@LMWH, bare OPNs were eliminated more rapidly by both healthy and colitis model mice (Fig. 2c, d). In the colitis model mice administered with OPNs@LMWH, we also observed a significantly greater particle accumulation in the colon, which was approximately 2-fold greater than that in colitis model mice given indocyanine green-labelled OPNs (ICG-OPNs) and 12-fold greater than that in healthy mice given ICG-OPNs or ICG-OPNs@LMWH (Fig. 2e, f). Notably, both OPNs and OPNs@LMWH were eliminated from the colon in a short time in healthy mice, while OPNs@LMWH were remained located in the inflamed colon extensively (Fig. 2e), demonstrating the excellent inflammation-targeting capability of OPNs@LMWH. These findings suggested that OPNs@LMWH possessed an inflammation-targeting effect in vivo. To further observe the distribution of the particles in detail, frozen sections of colon tissue were obtained 24 h after administering FA-OPNs or FA-OPNs@LMWH. We found that there was almost no residual fluorescence signal from the particles in normal mouse colon tissue and believed that the structurally intact intestinal barrier prevented the entry of the particles and successfully excreted them (Fig. 2g, Supplementary Fig. 11). A small amount of OPNs remained in the inflamed colon tissue. This could be attributed to nonspecific factors such as disruption of the intestinal barrier and abnormal colon function. More significantly, a substantial amount of OPNs@LMWH not only accumulated at the site of inflammation but also penetrated deeper into the submucosal layer heavily infiltrated by integrin αM+ inflammatory cells (Fig. 2g, Supplementary Fig. 11), which suggested the excellent integrin αM-targeting effect of LMWH.

### Suppression of ROS and proinflammatory cytokines by OPNs@LMWH

Since the precise targeting properties of OPNs@LMWH have been illustrated, we next evaluated the efficacy of OPNs@LMWH for immune regulation, including ROS scavenging and anti-inflammatory effects. First, a 24 h cytotoxicity test was performed to evaluate cell viability in vitro. The results indicated that OPNs@LMWH had the desired biocompatibility and negligible cytotoxicity at a concentration of 100 μg mL^-1^ with both macrophages and intestinal epithelial cells (Supplementary Fig. 12a-d). To investigate the ROS scavenging ability of OPNs@LMWH, RAW264.7 cells were stimulated with phorbol 12-myristate 13-acetate (PMA) after pretreatment with different nanoparticles. We observed significant concentration-dependent intracellular ROS scavenging behaviour by both the OPNs and OPNs@LMWH, while OPNs@LMWH exhibited greater scavenging capability due to their enhanced cellular uptake efficiency (Fig. 3a-d, Supplementary Fig. 13a-c). In a confirmatory experiment, we also found that LMWH had little anti-ROS activity (Supplementary Fig. 14a-d) and that the mass proportion of LMWH in OPNs@LMWH did not exceed 1 % (Supplementary Fig. 15), suggesting that the enhanced cellular uptake efficiency rather than the potential oxidative sensitivity of LMWH resulted in enhanced intracellular ROS scavenging behaviour. Then we evaluated the protective effects of the nanoparticles on peroxide-induced apoptosis in NCM460 cells, which represents the extracellular ROS scavenging behaviour and protection of the intestinal epithelium. We observed comparable results for OPNs and OPNs@LMWH (Fig. 3e, f), suggesting comparable extracellular ROS scavenging behaviour. Surprisingly, OPNs can also suppress the production of IL-6 and TNF-α, which are two essential proinflammatory cytokines that are involved in most autoimmune diseases. This effect was improved by the LMWH coating (Fig. 3g, h). Considering the mutually reinforcing relationship between oxidative stress and inflammation, we attribute this improvement in anti-inflammatory activity to increased cellular uptake and the consequent increase in intracellular ROS scavenging efficiency, where LMWH plays an important role. These results demonstrated that OPNs@LMWH are more effective at comprehensively reversing oxidative stress and inflammation.

**Fig. 3.**
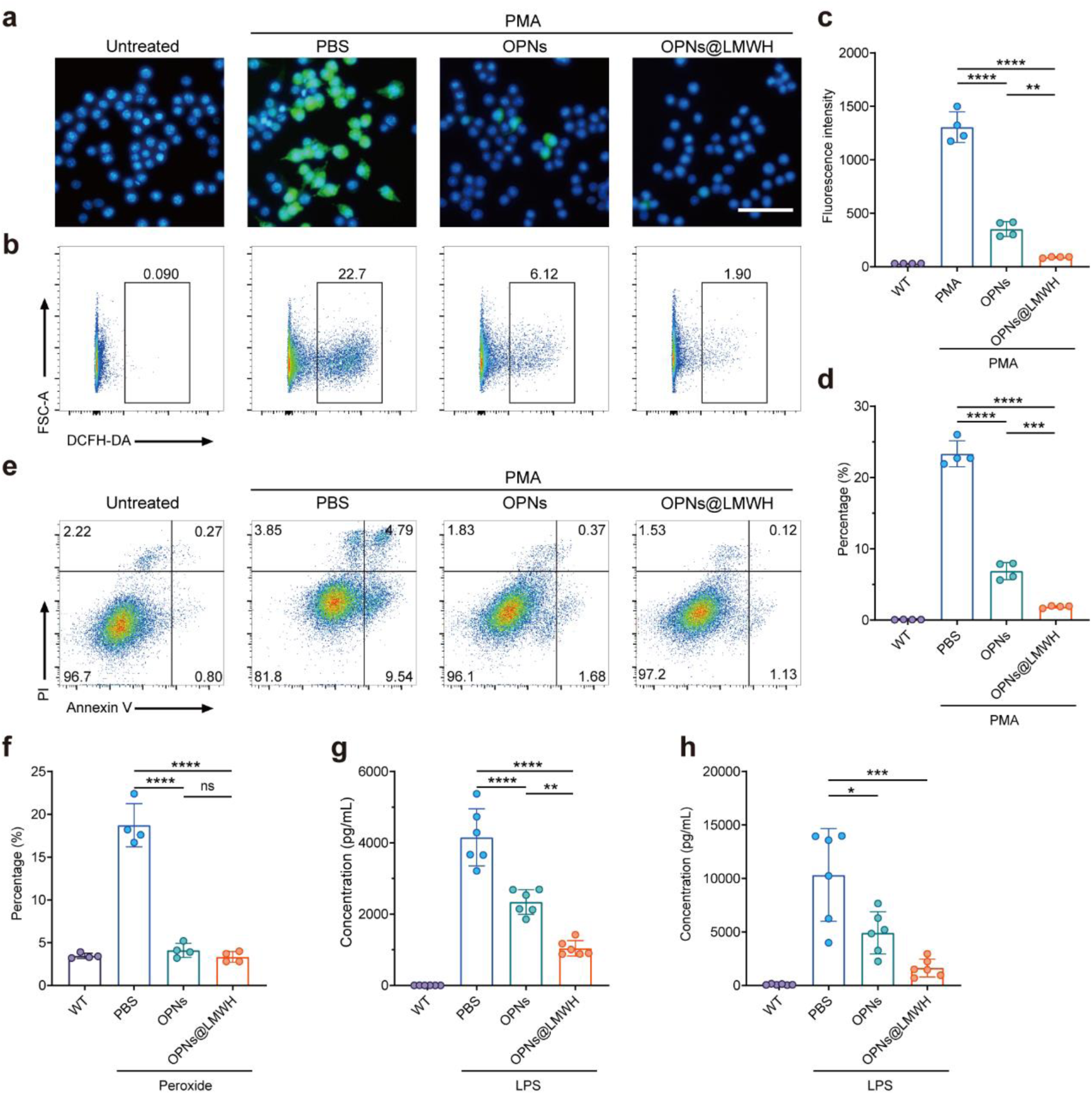
Suppression of ROS and proinflammatory cytokines by OPNs@LMWH. **a-d,** RAW264.7 cells were stimulated with PMA and treated with OPNs or OPNs@LMWH (10 μg mL^-1^). The level of intracellular ROS was detected by DCFH-DA, visualized by fluorescence microscopy (scale bar, 50 μm) (**a**) and determined by flow cytometry (**b**). The fluorescence intensity (**c**) and percentage of ROS-positive cells were calculated (**d**). **e, f,** NCM460 cells were incubated with H O (500 μM) and treated with OPNs or OPNs@LMWH (10 μg mL^-1^). Peroxide-induced apoptosis was detected by staining with Annexin-V and propidium iodide (PI) (**e**). The percentages of total apoptotic cells in the different treatment groups were calculated (**f**). **g, h,** RAW264.7 cells were stimulated with LPS to produce proinflammatory cytokines and incubated with OPNs or OPNs@LMWH (10 μg mL^-1^). The concentrations of IL-6 (**g**) and TNF-α (**h**) were determined via ELISA. Representative images, gating strategy and quantitative analysis of n=4 biologically independent samples (**a-f**) and n=6 biologically independent samples (**g, h**) are shown. The data are presented as the means ± SDs. * *p* < 0.05, ** *p* < 0.01, *** *p* < 0.001, **** *p* < 0.0001 and ns *p* > 0.05, analysed by ordinary one-way ANOVA (**c, d, f-h**) for multiple comparisons.

### Therapeutic efficacy of OPNs@LMWH administered orally for UC treatment

We next investigated the therapeutic efficacy of OPNs@LMWH in alleviating the manifestations of colitis in mice. We first examined the in vivo safety of OPNs@LMWH to confirm that a high daily dosage of OPNs@LMWH administered orally for two weeks did not cause significant damage to the mice. Our results indicated that there was no evidence-based damage in five main organs after constant daily oral administration at various concentrations, which preliminary suggested the ideal biosafety of the formulation (Supplementary Figs. 16-18). We subsequently investigated the therapeutic efficacy of OPNs and OPNs@LMWH in colitis model mice. Fig. 4a schematically shows the development of the DSS-induced colitis mouse model and the oral treatment strategies with OPNs or OPNs@LMWH. Compared with water, both OPNs and OPNs@LMWH significantly protected mice against colitis, including body weight loss (Fig. 4b), colon length shortening (Fig. 4c, e) and enlargement of the spleen (Fig. 4d, f). Importantly, compared with that of bare OPNs, a significant improvement was observed with OPNs@LMWH. According to the histopathological examination, treatment with OPNs@LMWH also protected the colonic epithelium from pathological damage, and the structures of the crypts were significantly more complete and more stable after OPNs@LMWH treatment (Fig. 4g, h). These results indicated that OPNs@LMWH could effectively alleviate disease status and that the LMWH coating could significantly improve the efficacy, which was consistent with the results of in vitro experiments and our speculation.

**Fig. 4.**
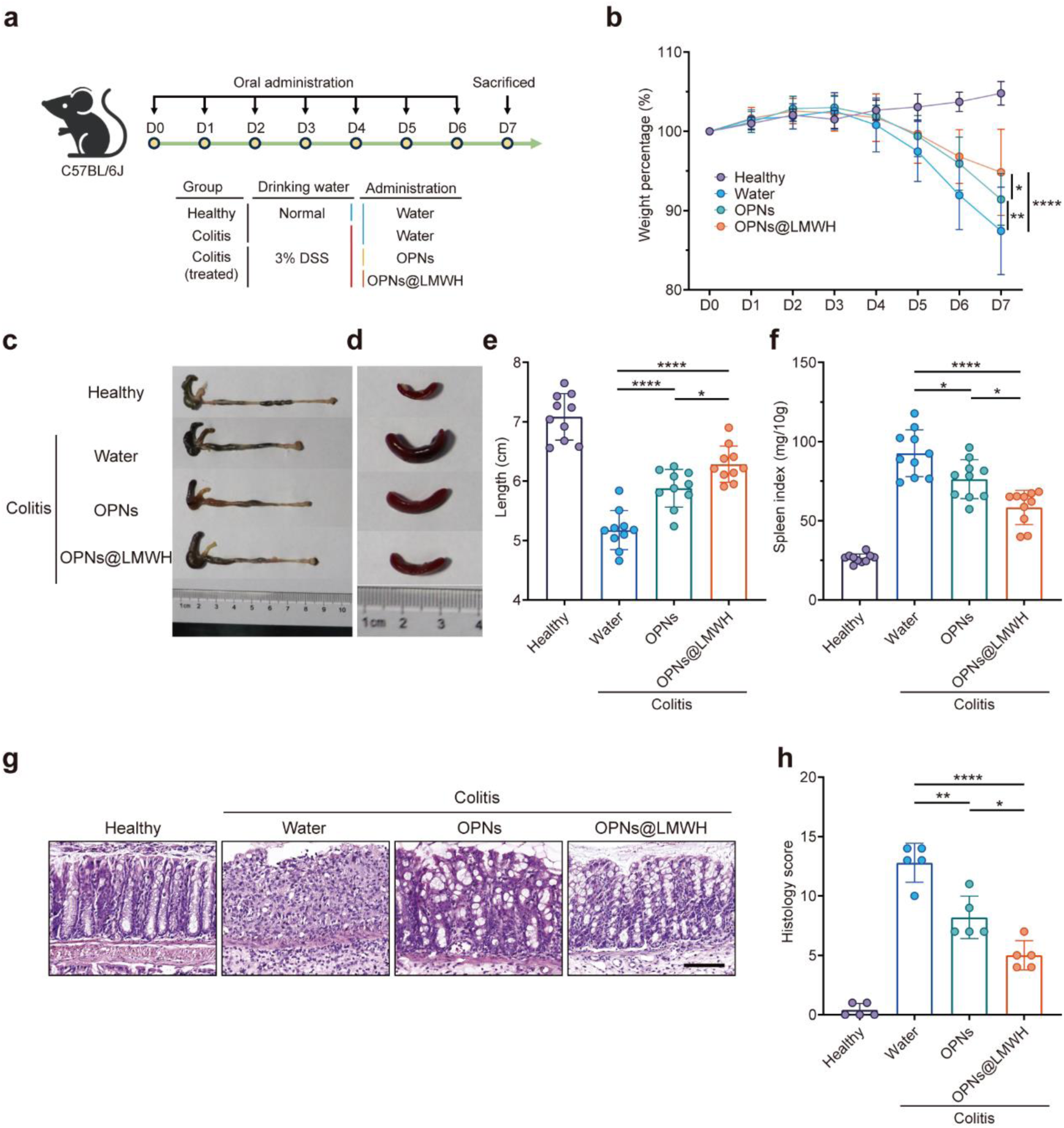
Evaluation of the therapeutic efficacy of OPNs@LMWH in mice with colitis. **a,** Schematic illustration of the development of the DSS-induced colitis mouse model. Mice were administered 50 mg kg^-1^ OPNs or OPNs@LMWH daily or in drinking water. **b,** Bodyweight changes in each group measured daily. **c-f,** Mice were sacrificed on day 7, and the colon (**c**) and spleen (**d**) were collected. The colon length (**e**) and spleen index (**f**) were quantified. **g,** Representative images of colons stained with haematoxylin and eosin. Scale bar, 100 μm. **h,** The histological score was calculated to quantify tissue damage. Representative images or quantitative analyses of n= 10 animals (**b-f**) and n=5 biologically independent samples (**g, h**) are shown. The data are presented as the means ± SDs. * *p* < 0.05, ** *p* < 0.01, *** *p* < 0.001, **** *p* < 0.0001 and ns *p* > 0.05, analysed by two-way ANOVA (**b**) or ordinary one-way ANOVA (**e, f, h**) for multiple comparisons.

### OPNs@LMWH restore intestinal homeostasis and protect the colonic epithelium

Our results described above revealed the potential of OPNs@LMWH for restoring intestinal redox homeostasis. Inspired by these data, we examined the mechanisms of OPNs@LMWH in UC management. Our enzyme-linked immunosorbent assay (ELISA) results confirmed that there were lower levels of proinflammatory cytokines in both the peripheral blood and lamina propria of colitis model mice treated with OPNs@LMWH than in those treated with pure water or OPNs (Fig. 5a-d). These results revealed the enhanced immune regulation ability of OPNs@LMWH compared with OPNs. According to the myeloperoxidase (MPO) immunohistochemistry (IHC) results, we observed the least oxidative stress in the colons of the colitis model mice in the OPNs@LMWH treatment group compared with the other groups. Remarkably, the decrease in the proportion of MPO-positive cells in the OPNs@LMWH treatment group was primarily attributed to the significant reduction in the submucosal layer, which is rich in recruited proinflammatory immune cells (Fig. 5e). These results indicated that OPNs@LMWH precisely influenced the production of excess proinflammatory cytokines in immune cells by scavenging ROS, which is consistent with our in vitro results. Moreover, the IHC results showed decreased activation of the JAK-STAT signalling pathway in the lamina propria after OPNs@LMWH treatment (Fig. 5f). As an essential signal transduction pathway, JAK-STAT is activated mainly by proinflammatory cytokines in colitis lesions, hence, decreasing activation of this pathway indicates a reduction in overall inflammatory activity.

**Fig. 5.**
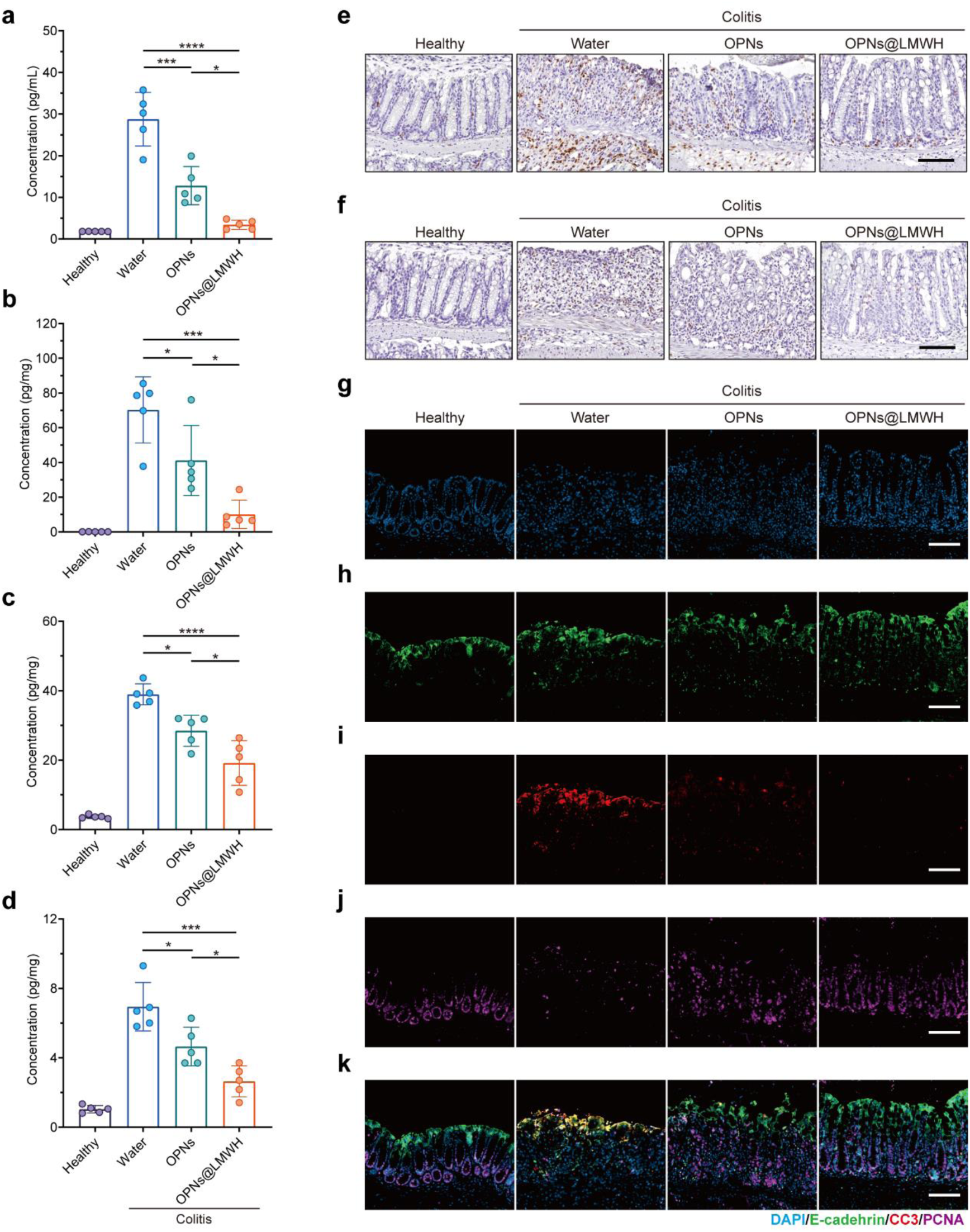
OPNs@LMWH restore intestinal homeostasis and protect the colonic epithelium. **a-d,** Colitis model mice were treated with OPNs or OPNs@LMWH. The concentrations of IL-6 in the serum (**a**) and IL-6 (**b**), TNF-α (**c**) and IL-1β (**d**) in colon tissue were determined via ELISAs. **e, f,** Representative images of immunohistochemical staining for MPO (**e**) and p-STAT3 (**f**). Scale bar, 100 μm. **g-k,** Representative images of immunofluorescence staining for proliferation and apoptosis showing DAPI (**g**), E-cadherin (**h**), PCNA (**i**), CC3 (**j**), and merged images (**k**) obtained via CLSM. Scale bar, 100 μm. Quantitative analysis or representative images from n=5 biologically independent samples (**a-d**) and n= 3 biologically independent samples (**e-k**) are shown. The data are presented as the means ± SDs. * *p* < 0.05, ** *p* < 0.01, *** *p* < 0.001, **** *p* < 0.0001 and ns *p* > 0.05, analysed by ordinary one-way ANOVA (**a-d**) for multiple comparisons.

An important sign of the restoration of intestinal homeostasis is the normalization of crypt maintenance and self-renewal. The stability and rapid renewal of the intestinal mucosal layer depend on the orderly arrangement of crypt cells and the performance of each functional cell, including maintaining the epithelial barrier and regulating cell proliferation and apoptosis^42^. Interestingly, by immunofluorescence staining, we found that the epithelial cells were restored and arranged more regularly in colitis model mice treated with OPNs@LMWH, which also indicated that intestinal mucosal permeability was restored (Fig. 5g, h). We utilized cleaved caspase 3 (CC3), a marker of apoptosis, to investigate the occurrence of apoptosis in colitis model mice. Our findings revealed substantial overlap between cells exhibiting high expression of CC3 and E-cadherin in untreated colitis model mice (Fig. 5h, i, k). Concurrently, the proliferation (marked by PCNA) of transit amplifying (TA) cells and intestinal stem cells located at the base of the crypts was virtually halted (Fig. 5j). This finding suggested that the mucosal layer in colitis model mice had lost its barrier function and self-repair capabilities. However, for colitis model mice treated with OPNs or OPNs@LMWH, a significant reduction in the proportion of apoptotic epithelial cells was observed, demonstrating that the proliferation of crypt cells was restored (Fig. 5i-k). More importantly, most of the PCNA-positive cells were located at the bottom of the crypts after treatment with OPNs@LMWH. These results indicated orderly proliferation and differentiation were ensured (Fig. 5j). Taken together, these findings suggested that OPNs@LMWH can significantly ameliorate UC in vivo by restoring intestinal homeostasis and protecting the colonic epithelium.

## Discussion

The intricacies of UC present significant challenges for existing treatment methods. Currently available clinical drugs have notable limitations, and the off-target effects of oral medications are concerning^8,43^. To address these issues, numerous studies on precision drug delivery systems for UC have been conducted, yielding promising results^11,12^. Despite the varying approaches and outcomes described in previous research, there is consensus that designing new multifunctional targeted therapies for the immune response holds great potential^18^.

In our work, we considered the transport and infiltration phenomena of inflammatory cells at UC lesion sites and proposed a “two-birds-one-stone” precise nanotherapy based on integrin αM targeting and ROS scavenging. By adopting a modular integration approach, OsEPL was combined with LMWH through nanoprecipitation and layer-by-layer self-assembly to form core-shell nanoparticles. Given the imbalance in redox homeostasis at inflammatory sites, we opted for a drug-free approach using biocompatible ROS scavenging materials derived from modified EPL as a regulator of intestinal oxidative stress.

Despite the discovery of the heparin–integrin interaction mechanism in previous research, our knowledge regarding its potential in drug delivery remains sparse. Our findings robustly validated the enhanced immune cellular uptake of heparin-coated nanoparticles and highlighted the crucial mediating role of LMWH-integrin αM interactions. OPNs@LMWH also demonstrated a significantly extended retention time in the colon of colitis model mice. Remarkably, 24 h after oral administration, the concentration of OPNs@LMWH was approximately double that of the bare OPNs. Conversely, normal mice rapidly eliminated OPNs@LMWH, underscoring its superior inflammation-targeting properties. A closer examination of colon tissue sections revealed that OPNs@LMWH penetrated more deeply into submucosal tissues with inflammatory cell infiltration than bare OPNs did. As anticipated, OPNs@LMWH successfully targeted intestinal inflammatory cell migration, accumulation, and infiltration in colitis model mice.

To understand how OPNs@LMWH improve colitis, we conducted comprehensive in vivo and in vitro experiments. In our in vitro studies, we noted a reduction in the levels of intracellular ROS and in the secretion of proinflammatory cytokines following the application of OPNs@LMWH. Interestingly, in experiments involving cells damaged by exogenous peroxides, the performance of OPNs@LMWH was comparable to that of OPNs. These findings suggest that while the cellular uptake efficiency of OPNs@LMWH is better than that of OPNs, their inherent antioxidant functionality remains effective. In subsequent in vivo trials, OPNs@LMWH demonstrated superior effectiveness in alleviating the symptoms of colitis. Our results revealed that following treatment with OPNs@LMWH, there were significant reductions in the levels of proinflammatory cytokines in peripheral blood and colonic lamina propria in mice with colitis. Additionally, we noted a marked decrease in oxidative stress within immune cells and reduced activation of the JAK-STAT signalling pathway. The process of apoptosis in the intestinal epithelial cells of the mice was effectively suppressed, and their orderly proliferation at the base of the crypts was successfully reinstated.

Our findings provide robust validation for our “two-birds-one-stone” strategy that combines LMWH-based integrin targeting with the regulation of intestinal redox homeostasis, thereby highlighting the application of OPNs@LMWH in the treatment of UC. More promisingly, our research underscores the feasibility and flexibility of employing LMWH via oral administration as an integrin-targeted treatment for UC. We postulate that the LMWH surface coating could be universally implemented across a variety of nanoparticle designs targeting integrins αM. This straightforward and adaptable modular design could stimulate further research into integrin-targeted treatment strategies for autoimmune diseases.

## Experimental section

### Materials

ε-Polylysine (EPL), 4-hydroxyphenyl boronic acid pinacol ester, dichloromethane (DCM), carbonyl diimidazole (CDI), sodium chloride (NaCl), sodium sulfate (Na_2_SO_4_), 4-dimethylaminopyridine (DMAP), dimethyl sulfoxide (DMSO), lecithin, ammonium acetate (CH_3_COONH_4_) and hydrogen peroxide solution (H_2_O_2_) were purchased from Aladdin (Shanghai, CHN). Heparin was purchased from Changshan Biotechnology Co., Ltd. (Hebei, CHN). Chitosan (CS, >75 % deacetylated), indocyanine green (ICG), fluorescein amine (FA), azure A, lipopolysaccharide (LPS), phorbol 12-myristate 13-acetate (PMA), sucrose and haematoxylin solution were purchased from Sigma–Aldrich (St. Louis, MO, USA). Cell Counting Kit-8 (CCK-8), a BCA protein assay kit, protease and phosphatase inhibitor cocktail for mammalian cell and tissue extracts (P1050) and DCFH-DA were purchased from Beyotime Biotechnology (Shanghai, CHN). Foetal bovine serum (FBS) was purchased from Gemini (USA). Dulbecco’s modified Eagle’s medium (DMEM), Roswell Park Memorial Institute (RPMI) 1640 medium, and Opti-MEM I reduced serum medium were purchased from Gibco (USA). Dextran sulfate sodium (DSS, colitis grade) was purchased from MP Biomedicals (USA). The anti-E-cadherin antibody (14-3249-82), CM-DiI (C7001), and the ELISA kits for IL-6 (88-7064), TNF-α (88-7324), and IL-1β (88-7013) were purchased from Invitrogen (USA). The Annexin-V/PI kit was purchased from BioLegend, Inc. (USA). The antibodies against phospho-STAT3 (Tyr705, 9145), cleaved caspase-3 (Asp175, 9579) and PCNA (2586) were purchased from Cell Signaling Technology (USA). The antibodies against myeloperoxidase (ab208670) and integrin αM (ab133357) were purchased from Abcam (Cambridge, UK). DAPI was purchased from Thermo Scientific (USA). Secondary antibodies conjugated with Alexa Fluor 488 (A-21208), Alexa Fluor 594 (A-21207), and Alexa Fluor 647 (A-31573 and A-31571) were purchased from Invitrogen (USA). Secondary antibodies conjugated with horseradish peroxidase (HRP), 3,3’-diaminobenzidine (DAB), peroxidase blocking solution and serum-free protein blocking solution were purchased from Dako (DNK). The O.C.T. compound was purchased from Sakura (USA).

### Synthesis of OsEPL

Oxidation-sensitive ε-polylysine (OsEPL) was synthesized inspired by Broaders et al.^44^. Briefly, 4-hydroxyphenyl boronic acid pinacol ester (10 mmol) was dissolved in DCM (10 mL). Then, CDI (20 mmol) was added, and the mixture was stirred at RT for 1 h. The solution was diluted by adding another 10 mL of DCM and washed with dd-H_2_O (20 mL) three times. The obtained organic phase was subsequently washed with saturated brine (20 mL) three times, dried over anhydrous Na_2_SO_4_ for 30 min and concentrated in a rotary evaporator to obtain imidazoyl carbamate (Compound 1, 8.5 mM, 85 %). To generate OsEPL, Compound 1 (820 mg), EPL (320 mg) and DMAP (305 mg) were dissolved in anhydrous DMSO (10 mL). The mixture was stirred at RT for 24 h. Then, OsEPL was precipitated in dd-H_2_O (40 mL) and isolated by centrifugation (8000 × g, 10 min). The precipitate was washed with dd-H_2_O (20 mL) three times and lyophilized (72 h) for further use.

### Preparation of OPNs@LMWH

LMWH (Mw=8.5 kDa) was prepared by a controlled heparinase depolymerization method according to previous methods.^26,45^ The molecular weights of unfractionated heparin (UFH) and LMWH were determined by high-performance liquid chromatography (HPLC, 1260 Infinity II, Agilent, USA) with a GPC column (TSK Gel G2000SWXL, 7.8 mm × 300 mm, Tosoh Co., JPN), and the results are shown in Supplementary Fig. 6. OPNs@LMWH were prepared by a nanoprecipitation/layer-by-layer method. Briefly, lecithin (20 mg) was dispersed in 1 mL of anhydrous ethanol and subsequently added dropwise to 8 mL of CH_3_COONH_4_ buffer (0.1 M, pH = 5.00) with magnetic stirring (600 rpm) to form a dispersion. OsEPL (20 mg) was dissolved in DMSO (1 mL) and subsequently added dropwise to the dispersion. The nanoparticles were collected by centrifugation (30000 × g, 10 min). Bare OPNs were harvested after washing three times with dd-H_2_O. CS (20 mg) was dissolved in 10 mL of CH_3_COONH_4_ buffer (0.1 M, pH = 5.00). Then, the bare OPNs were dispersed in the CS solution and incubated at RT for 30 min with magnetic stirring (600 rpm). The CS-coated OPNs were collected after washing three times with dd-H_2_O, dispersed in 10 mL of LMWH aqueous solution (2 mg mL^-1^) and incubated at RT for 30 min with magnetic stirring (600 rpm) again. Finally, the OPNs@LMWH were washed three additional times and collected by centrifugation (30000 × g, 10 min) for further use. For in vivo imaging, ICG (1 mg mL^-1^) or FA (1 mg mL^-1^) was added to the primary DMSO solution containing OsEPL, and two kinds of fluorescently labelled OPNs or OPNs@LMWH were prepared via a method same to that described above. The fluorescence intensity of the fluorescently labelled nanoparticles was normalized before use.

### Characterization of OPNs@LMWH

The surface morphologies of the OPNs and OPNs@LMWH were determined via scanning electron microscopy (SEM) (Merlin, Zeiss, DEU), transmission electron microscopy (TEM) (Tecnai G2 20, FEI, USA) and aberration-corrected transmission electron microscopy (AC-TEM) (Titan 80-300, FEI, USA). The sizes, polydispersity index (PDI) and surface charges of the OPNs and OPNs@LMWH were analysed via dynamic light scattering (DLS) (Zetasizer Nano ZS90, Malvern, UK). For stability assays, simulated gastric fluid (SGF) and simulated intestinal fluid (SIF) were prepared according to the methods of the United States Pharmacopeia. For stability assay, the nanoparticles were dispersed and incubated in deionized water (48 h) at 4 °C or in SGF (2 h) or SIF (44 h) at 37 °C. Degradation was evaluated by measuring the absorbance at 500 nm with a microplate reader (Infinite M200 Pro, Tecan Trading AG, Switzerland). The mass proportion of LMWH in OPNs@LMWH was calculated by measuring the concentration of free LMWH in the supernatant before and after coating. The concentration of LMWH was detected by incubation with Azure A.

### Cell culture

RAW264.7 (mouse) macrophages and NCM460 (human) colonic enterocytes were kindly provided by Prof. Zhijie Chang (School of Medicine, Tsinghua University). RAW264.7 cells were cultured in DMEM supplemented with 10 % FBS, and NCM460 cells were cultured in RPMI 1640 medium supplemented with 10 % FBS. For in vitro assays, the medium was replaced with Opti-MEM I reduced serum medium after the cells were seeded overnight to avoid potential impacts from FBS.

### In vitro cellular uptake assays

RAW264.7 cells were seeded in 12-well plates (Corning 3513, USA) at a density of 3 × 10^5^ cells per well and stimulated with LPS (100 ng mL^-1^) and IFN-γ (20 ng mL^-1^) for 12 h to form M1 macrophages. Then, FA-labelled nanoparticles (20 μg mL^-1^) were added. After 8 h of incubation, the cells were collected and washed with PBS three times. The fluorescence intensity of the cells was immediately assessed via flow cytometry (LSRFortessa, BD, USA). To confirm the improved cellular uptake efficiency, free FA with the same fluorescence intensity as the nanoparticles was used as a control group. For immunofluorescence (IF) imaging, cells were seeded on round coverslips and pretreated with anti-integrin αM antibodies for 1 h before 8 h of incubation with the nanoparticles. Cells on coverslips were fixed in 4 % paraformaldehyde for 15 min, stained with CM-Dil and DAPI and washed three times. Confocal laser scanning microscopy (CLSM; FV3000RS, Olympus, JPN) was used to obtain images.

### In vitro ROS scavenging behaviour and anti-inflammatory effects

Intracellular ROS levels were evaluated via the DCFH-DA method. Briefly, RAW264.7 cells were seeded in 12-well plates (Corning 3513, USA) at a density of 3 × 10^5^ cells per well. After incubation for 12 h, the medium was replaced with Opti-MEM containing nanoparticles at a predetermined concentration, and the mixture was incubated for another 12 h. Then, PMA (100 ng mL^-1^) and DCFH-DA were added. After incubation for 1 h, the cells were collected and washed with PBS three times. The fluorescence intensity of the cells was immediately assessed by flow cytometry. To confirm the ROS scavenging function of the nanoparticles, we evaluated the reduction in peroxide-induced apoptosis via the Annexin-V/PI method. Briefly, NCM460 cells were seeded in 12-well plates (Corning 3513, USA) at a density of 3 × 10^5^ cells per well. After incubation for 12 h, the medium was replaced with Opti-MEM containing nanoparticles (10 μg mL^-1^), and the cells were incubated for 1 h. Then, H O (500 μM) was added, and the plate was incubated for 12 h. The cells were collected, stained with an Annexin-V/PI apoptosis kit and assessed immediately by flow cytometry. For the in vitro anti-inflammatory assay, RAW264.7 cells were seeded in 96-well plates (Corning 3599, USA) at a density of 3 × 10^4^ cells per well. After incubation for 12 h, the medium was replaced with Opti-MEM containing LPS (100 ng mL^-1^). After 30 min of stimulation with LPS, nanoparticles at a predetermined concentration (10 μg mL^-1^) were added, and the cells were incubated for an additional 24 h. The supernatant was collected to determine the concentration of proinflammatory cytokines by ELISA kits.

### Animals and colitis model

All animal studies complied with the regulations of the Laboratory Animal Resources Center, Tsinghua University (23-XXH2). Six-week-old male C57BL/6J mice (Specific Pathogen Free, SPF) were obtained from Charles River (Beijing, CHN) and maintained on a 12-h light–dark cycle at 25 °C in SPF barrier environment. Mice supplied with food and water were cohoused for 7 days before random assignment to the experimental groups. A colitis model was established by providing drinking water containing 3 % DSS, and this drinking water was renewed every other day.

### In vivo therapeutic efficacy

Mice were orally administered drinking water or nanoparticle dispersions at a dose of 50 mg kg^-1^ every day for 7 days (beginning on day 0 and ending on day 6). On day 7, the mice were sacrificed, and the spleens, colons and serum were excised. To obtain histological sections, colons were fixed in 4 % paraformaldehyde for 24 h, dehydrated and embedded in paraffin. The samples were cut into 5 μm thick sections by using a microtome (RM2235, Leica, DEU) and stored at −20 °C until further use. Haematoxylin and eosin staining was performed for histopathological examination. The histological scores were determined according to a previously reported method^46^. Briefly, infiltration of inflammatory cells (0: none, 1: mild, 2: moderate, 3: severe), depth of ulceration involvement (0: none, 1: mucosa, 2: submucosa, 3: muscular layer), area of ulceration involvement (0: none, 1: 0 %–25 %, 2: 26 %–50 %, 3: 51 %–75 %, 4: 76 %–100 %), and crypt destruction (0: none, 1: lower than one third, 2: lower than two thirds, 3: more than two thirds, 4: all crypts and epithelium) were assessed. To determine the concentrations of cytokines in colon tissues, colon segments were homogenized (tissue homogenizer, 30 Hz, 2 min) in PBS containing 2 % protease and phosphatase inhibitor cocktail. The supernatants were collected, and cytokine levels were assessed via ELISA kits. All samples were normalized to equivalent protein concentrations by using a BCA protein assay kit.

### Immunohistochemistry and immunofluorescence imaging of histological sections

Immunohistochemistry (IHC) and IF were used to investigate the underlying mechanisms. Briefly, histological sections were obtained by the same method as described for “In vivo therapeutic efficacy”, followed by deparaffinization and hydration of tissue sections through xylene and gradient ethanol treatment, respectively. Antigen retrieval was performed at 100 °C for 15 min in Tris-EDTA buffer (pH = 9.00). For IHC, a peroxidase blocking solution was applied to eliminate nonspecific staining, while for IF, a serum-free protein blocking solution was applied. The tissue slides were incubated overnight at 4 °C with primary antibodies. All the slides were washed with PBS (containing 0.05 % Tween-20) and incubated with secondary antibodies for 30 min at RT, where antibodies conjugated with HRP were used for IHC and antibodies conjugated with fluorophores were used for IF. For IHC, DAB was applied to develop the colour, followed by haematoxylin staining. For IF, nuclear staining was performed by using DAPI. Then, all the slides were mounted with mounting medium (Leica), protected from light and stored at 4 °C.

### In vivo biodistribution

Both healthy mice and mice with colitis were orally administered fluorescently labelled OPNs/OPNs@LMWH. Briefly, after fasting overnight, mice orally administered ICG-OPNs/OPNs@LMWH were imaged by an IVIS (Lumina III, USA) at 6 h, 12 h and 24 h after gavage and then sacrificed for colon imaging. To investigate the specific distribution of the labelled OPNs/OPNs@LMWH in colon tissue, mice orally administered FA-OPNs/OPNs@LMWH were sacrificed 24 h after gavage. The colons were obtained and stored in PBS containing 30 % sucrose for 24 h. Then, the solution was removed, and all the samples were embedded in O.C.T. compound. Histological sections were obtained by using a freezing microtome (CM3050S, Leica, DEU). Slides were stained with DAPI, CM-Dil and integrin-αM, mounted, protected from light and stored at 4 °C.

### Safety assays

In vitro safety assays utilized cells. Briefly, cells were cultured in 96-well plates with different concentrations of particles for 24 h. Then, CCK-8 reagent was added, and the plates were incubated for another 30 min. The absorbance of each well at 450 nm was measured, and this value was used to calculate cell viability. In vivo safety assays were based on mouse data. Briefly, mice were orally administered different doses of nanoparticles daily for 14 days, and their body weights were recorded every day. Then, all the mice were sacrificed on day 14. Blood samples were collected and used for blood haematology examinations. The spleens, hearts, lungs, livers and kidneys of the mice were collected to obtain sections that were stained with haematoxylin and eosin.

### Statistical analysis

The data are expressed as the mean ± standard deviation (SD). Statistical analysis was performed by student’s t tests for unpaired data and by one-way ANOVA or two-way ANOVA for multiple comparisons. A two-tailed *p* value of < 0.05 was considered to indicate statistical significance, denoted as * *p* < 0.05, ** *p* < 0.01, *** *p* < 0.001, **** *p* < 0.0001 and ns (not significant) for *p* > 0.05. GraphPad Prism 8.3 software was used for all statistical analyses.

## Data availability

All the data supporting the results of this study are available within the paper and its Supplementary Information. Additional processed data are available from the corresponding authors upon request.

## Acknowledgements

The authors thank Prof. Zhibing Zhang, FREng, University of Birmingham, for his invaluable advice in the preparation of manuscript. The authors thank Prof. Zhijie Chang, Tsinghua University, for generously providing the cell lines. We thank the National Center for Electron Microscopy in Beijing, School of Materials Science, Tsinghua University, for assistance with electron microscopy imaging; the Laboratory Animal Resources Center, Tsinghua University, for technical support; and the Center of Biomedical Analysis, Tsinghua University, for assistance with imaging technology and flow cytometry. This work was supported by grants from the Key Research and Development Program of the Ministry of Science and Technology (2023YFA0914300), the Shenzhen Medical Research Fund (SMRF, B2302009) and the National Natural Science Foundation of China (22161142014).

## Author contributions

XH. X. Y. W. and CY. Z. oversaw the research. L. H., B. X., Y. W., CY. Z. and XH. X. designed the research strategy. L. H. synthesized the materials and fabricated the nanotherapy. L. H., LQ. H., W. H., QX. Z., ZY. S., and HY. T. performed the experiments. L. H. and Y. W. analysed the experimental data. L. H., Y. W., CY. Z. and XH. X. prepared the manuscript. All the authors approved the manuscript.

## Supplementary Information

**Supplementary Scheme 1.**
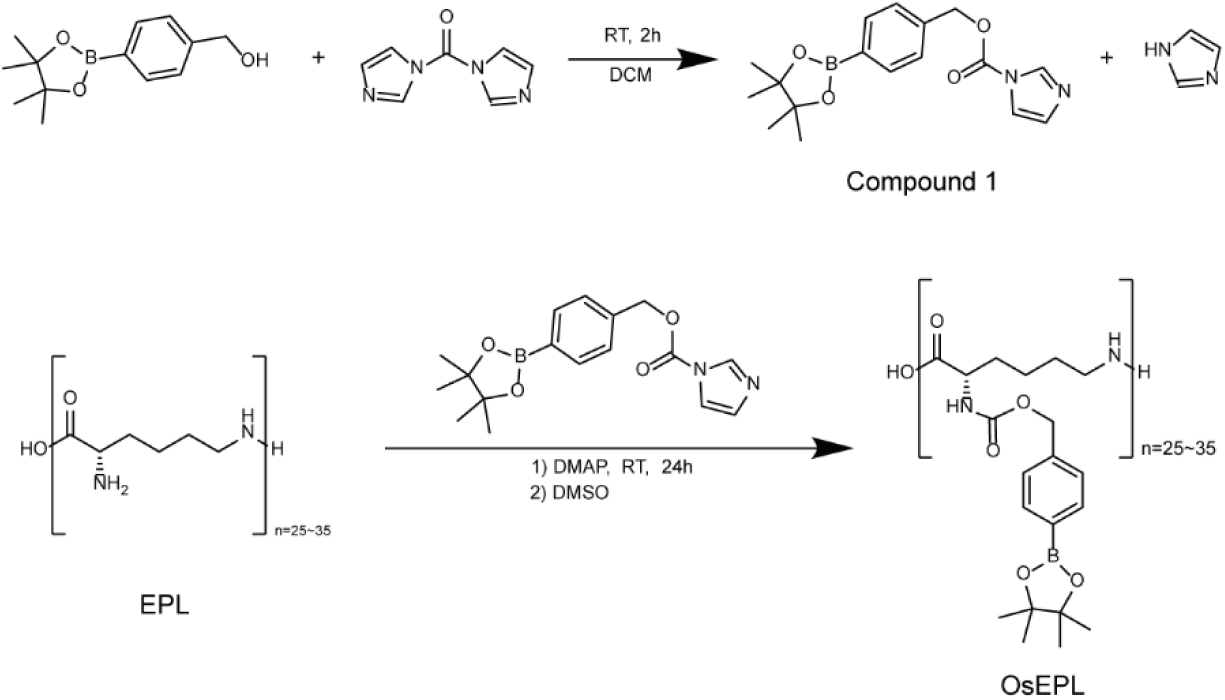
Synthesis of OsEPL.

**Supplementary Fig. 1.**
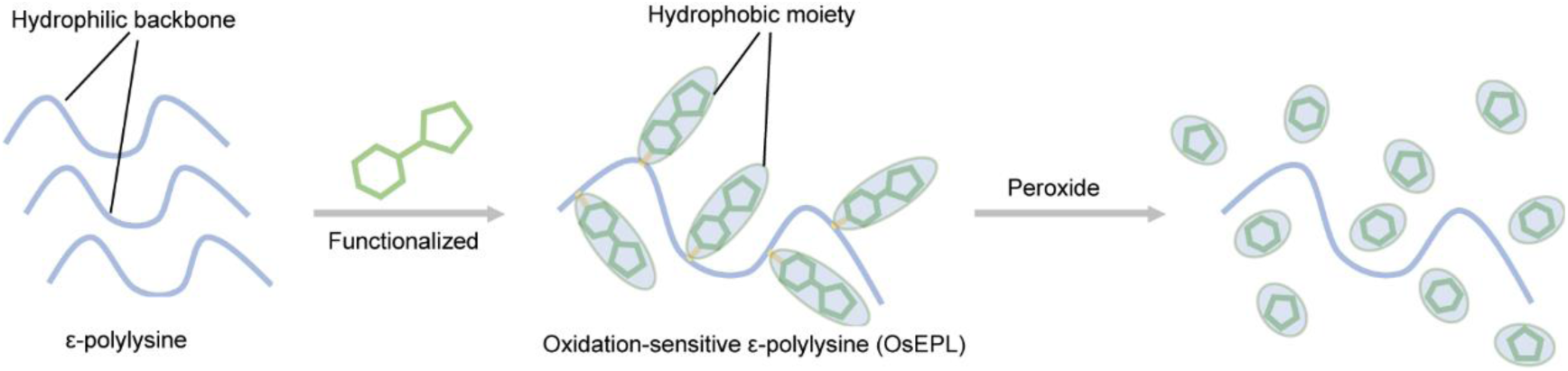
Schematic illustration of the preparation and degradation of OsEPL.

**Supplementary Fig. 2.**
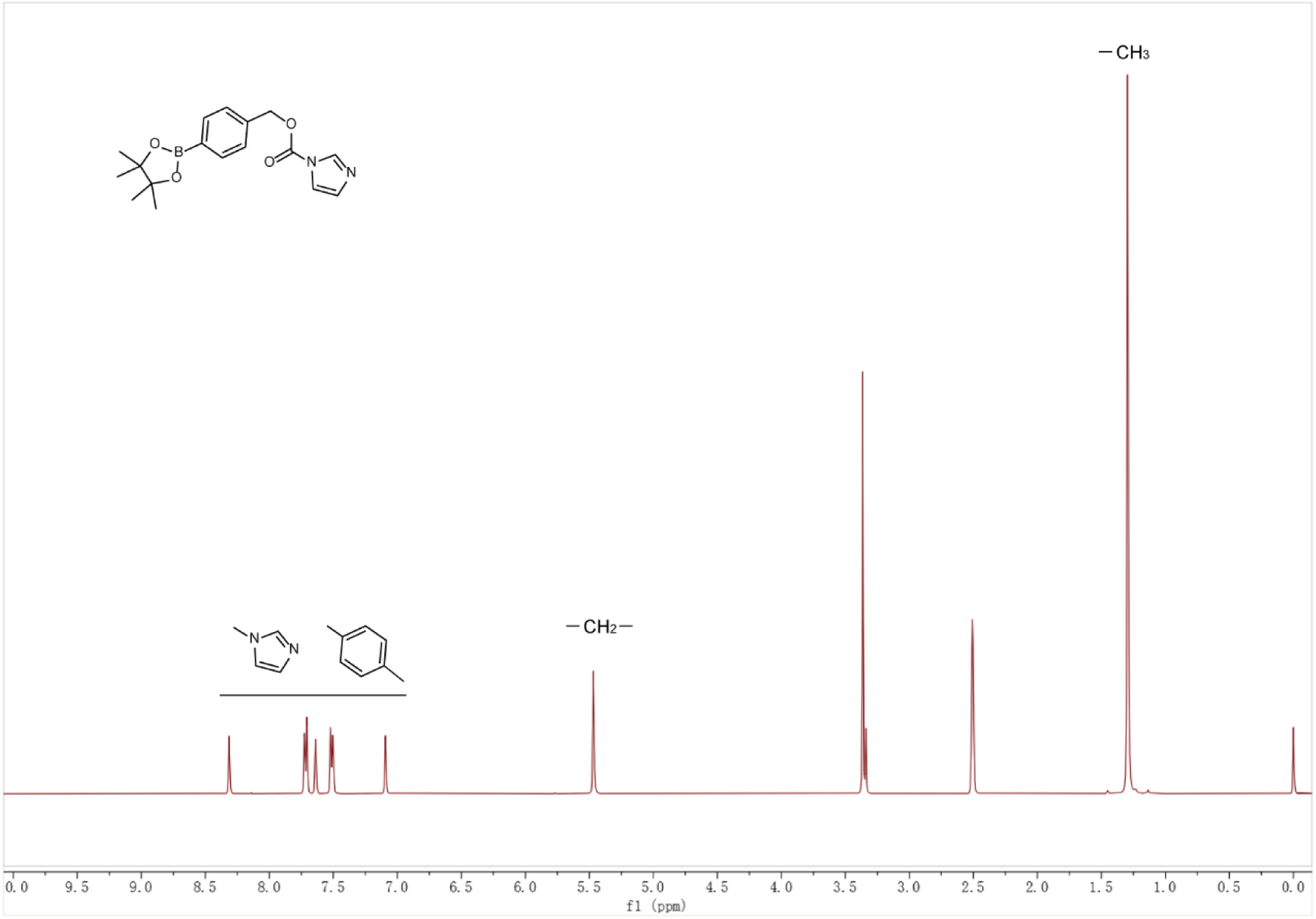
^1^H-NMR spectrum of imidazoyl carbamate (Compound 1) in DMSO-d_6_.

**Supplementary Fig. 3.**
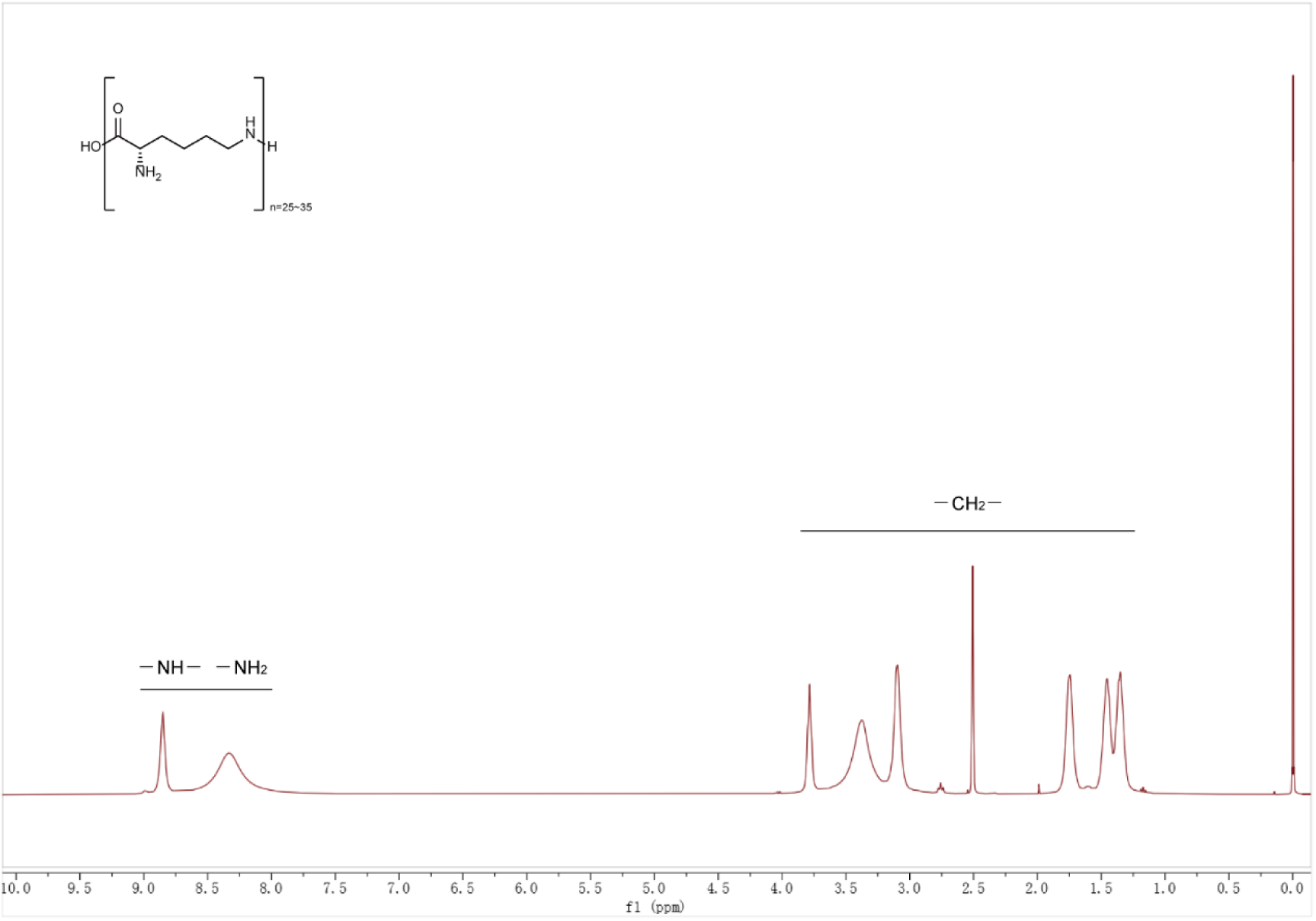
^1^H-NMR spectrum of EPL in DMSO-d_6_.

**Supplementary Fig. 4.**
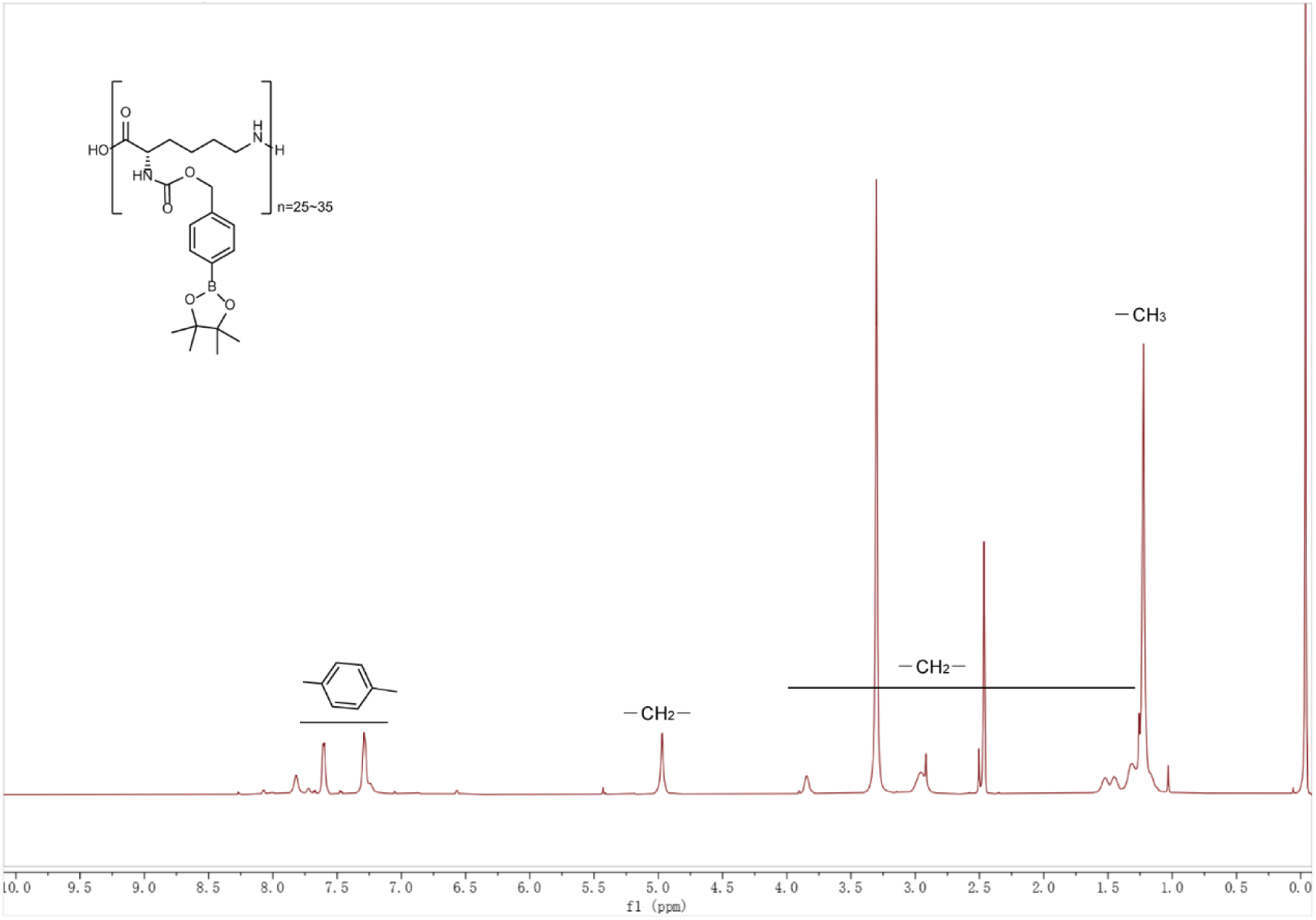
^1^H-NMR spectrum of OsEPL in DMSO-d_6_.

**Supplementary Fig. 5.**
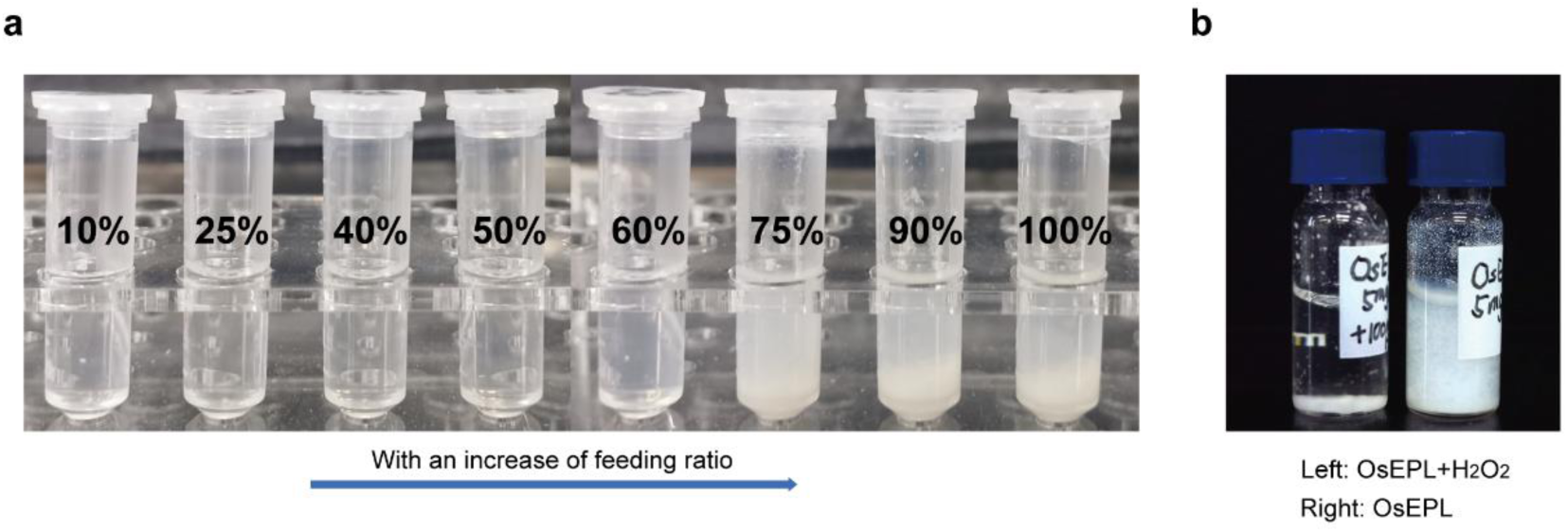
Physicochemical properties of OsEPL. **a,** A series of OsEPLs with different feeding ratios of phenylboronic acid pinacol ester and amino moieties presented varying amphiphilicities. **B,** OsEPL was incubated with 100 mM H_2_O_2_ for one hour.

**Supplementary Fig. 6.**
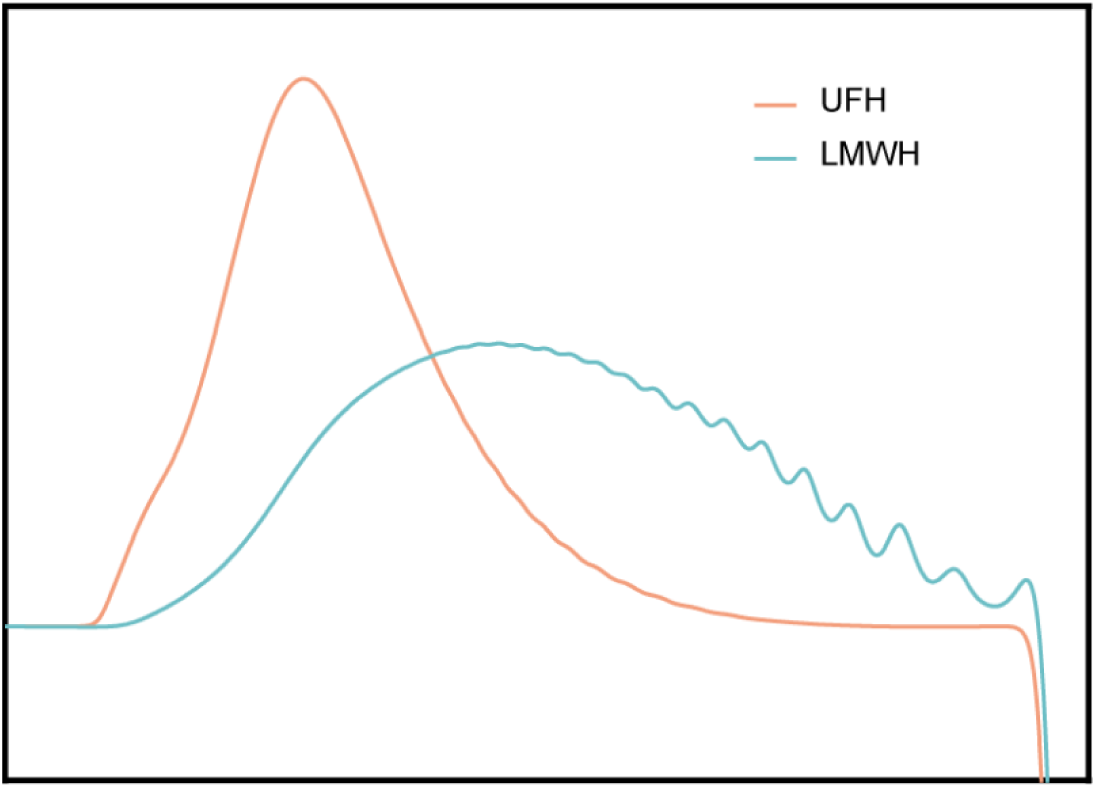
Molecular weight distributions of UFH and LMWH determined by GPC.

**Supplementary Fig. 7.**
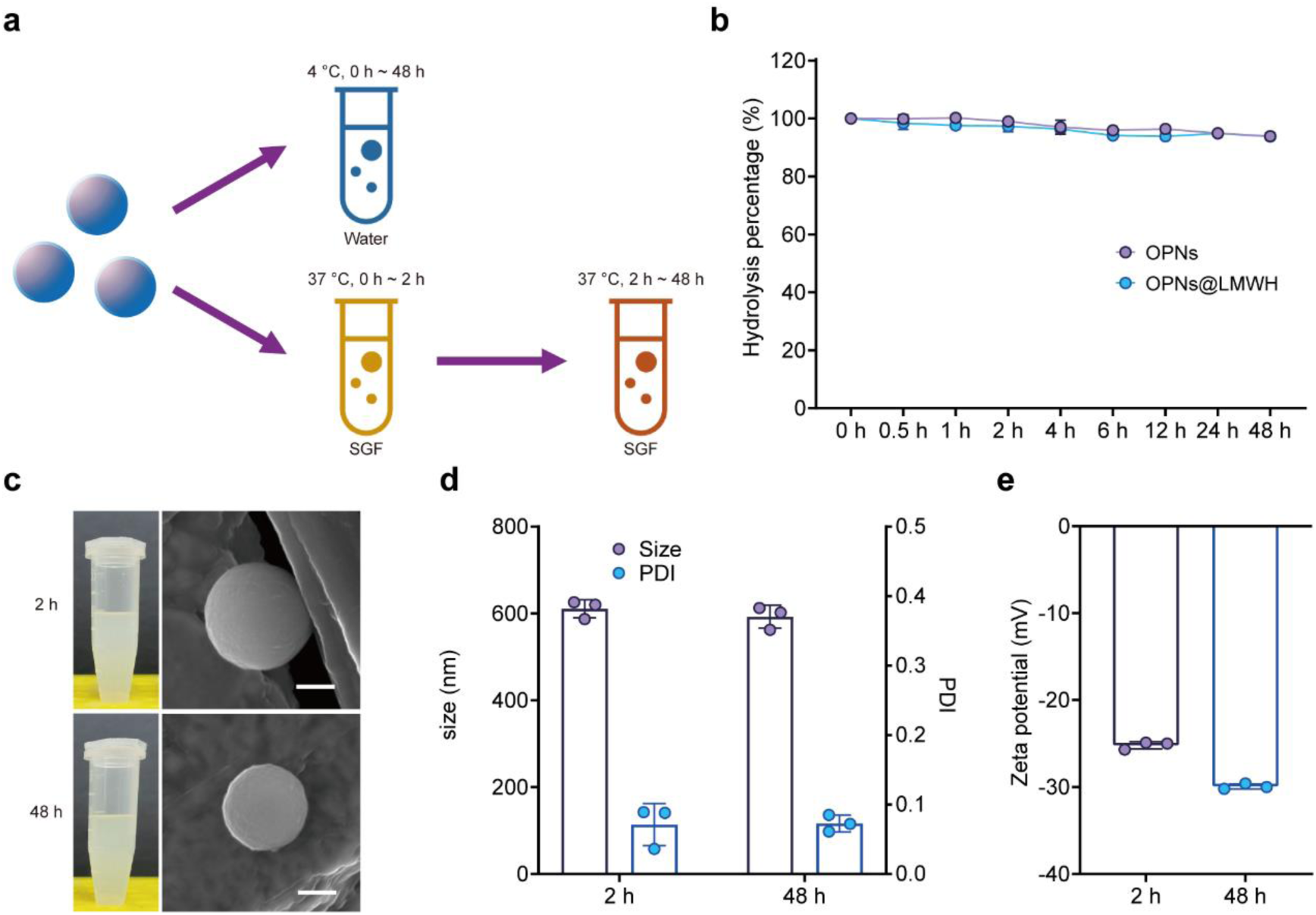
Stability of OPNs and OPNs@LMWH. **a,** OPNs and OPNs@LMWH were incubated under storage conditions (deionized water, 4 °C) or subjected to a simulated digestion process (SGF, 2 h, 37 °C; SIF, 46 h, 37 °C). **b,** The ratio of hydrolysed nanoparticles after storage. The data shown are from quantitative analyses of n=3 biologically independent samples. **c-e,** Representative SEM images (scale bar, 200 nm) (**c**), particle sizes (**d**) and zeta potentials (**e**) of OPNs@LMWH subjected to a simulated digestion process. The data are presented as the means ± SDs.

**Supplementary Fig. 8.**
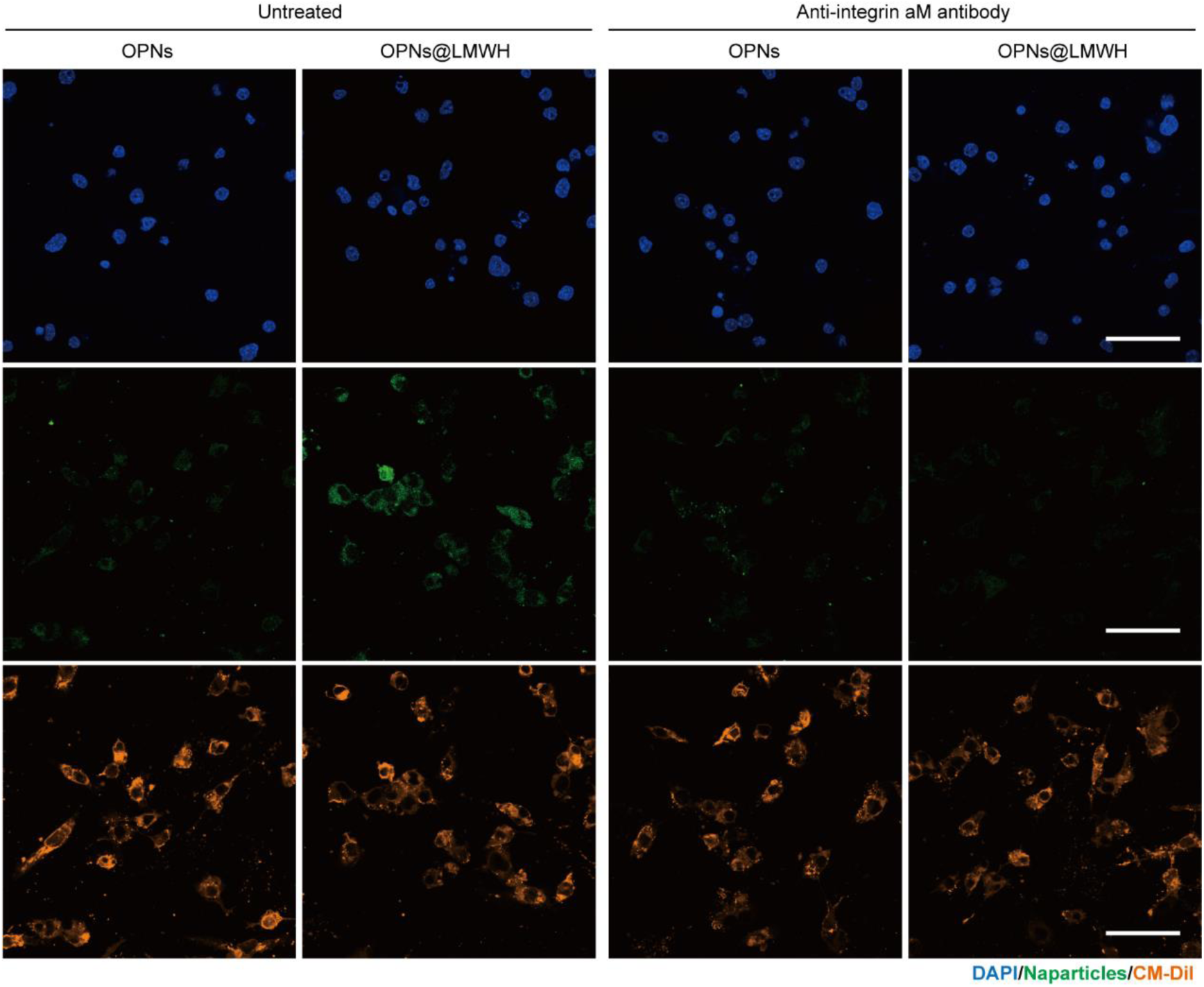
OPNs@LMWH internalization by M1 macrophages. M1-type RAW264.7 cells were incubated with FA-OPNs or FA-OPNs@LMWH after pretreatment with or without the anti-integrin αM antibody and visualized via CLSM. Scale bar, 50 μm. Representative images of n=6 biologically independent samples are shown.

**Supplementary Fig. 9.**
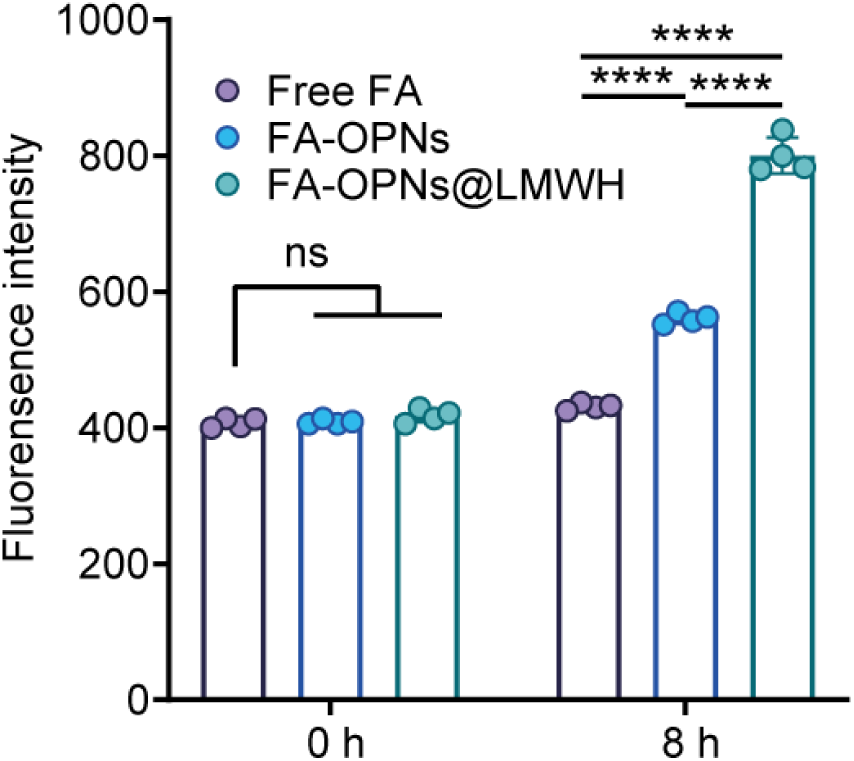
Quantitative analysis of the FA fluorescence intensity of the internalized nanoparticles. FA fluorescence intensity measurements by flow cytometry confirmed that free FA did not enter the cells. Quantitative analysis of 4 biologically independent samples is shown. The data are presented as the means ± SDs. * *p* < 0.05, ** *p* < 0.01, *** *p* < 0.001, **** *p* < 0.0001 and ns *p* > 0.05, analysed by two-way ANOVA multiple comparisons test.

**Supplementary Fig. 10.**
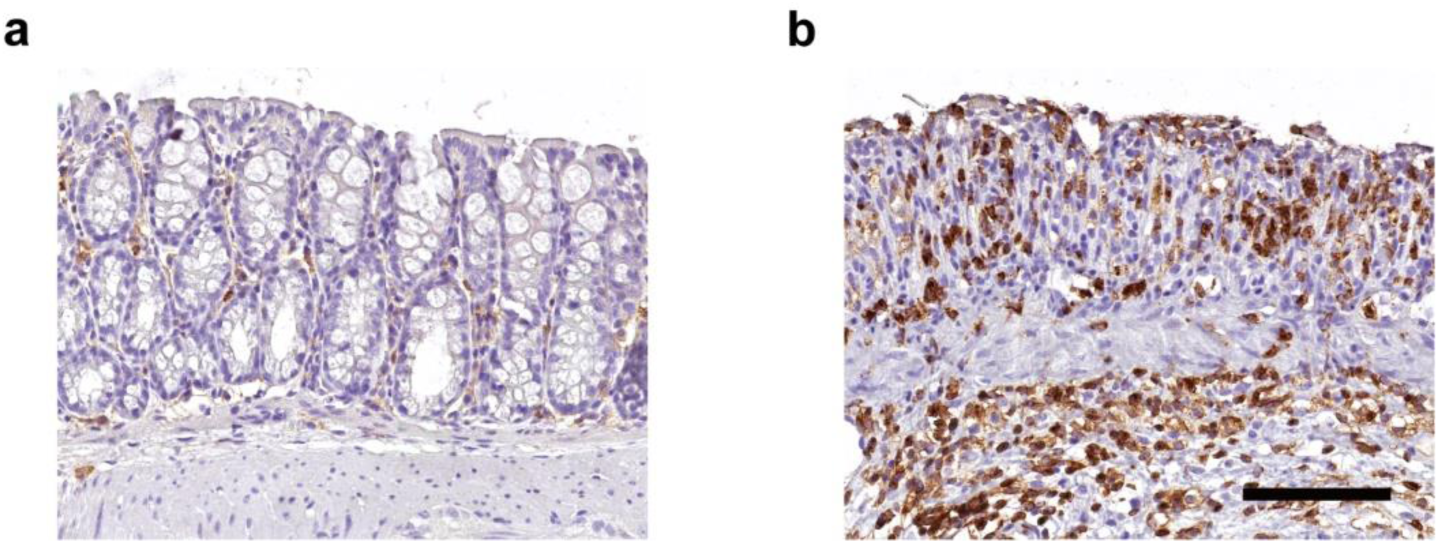
The high level of CD11b in the colonic submucosal layer of colitis model mice. Representative images of immunohistochemical staining for CD11b in healthy (**a**) or colitis (**b**) mice. Scale bar, 100 μm. Representative images from n= 3 biologically independent samples are shown.

**Supplementary Fig. 11.**
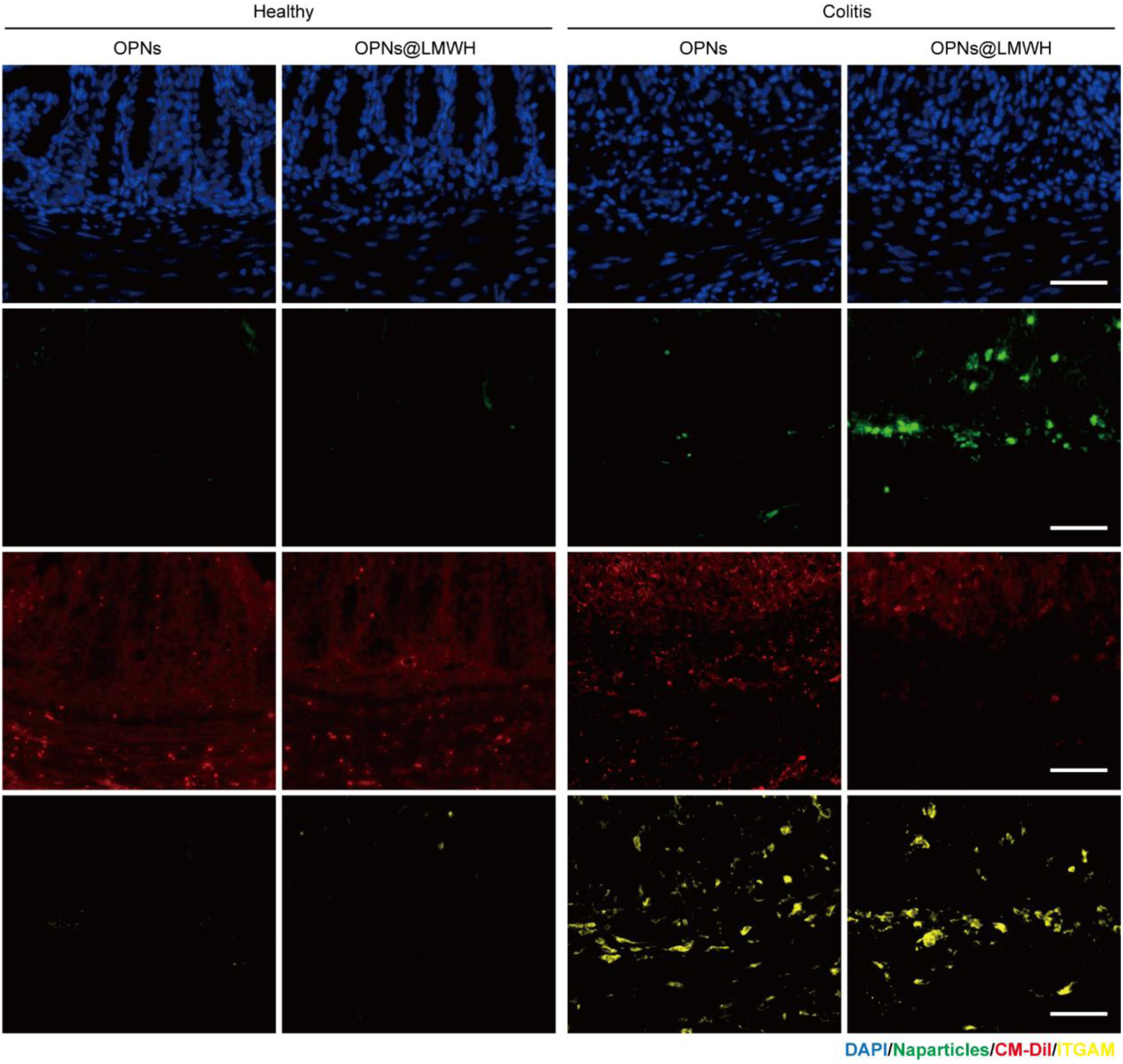
OPNs@LMWH accumulate in inflamed colon tissue infiltrated by inflammatory cells. Healthy and colitis model mice were orally administered FA-OPNs or FA-OPNs@LMWH, and colon tissues were collected after 24 h and stained with DAPI, CM-Dil and integrin-αM and visualized via CLSM. Scale bar, 50 μm. Representative images of n=3 animals are shown.

**Supplementary Fig. 12.**
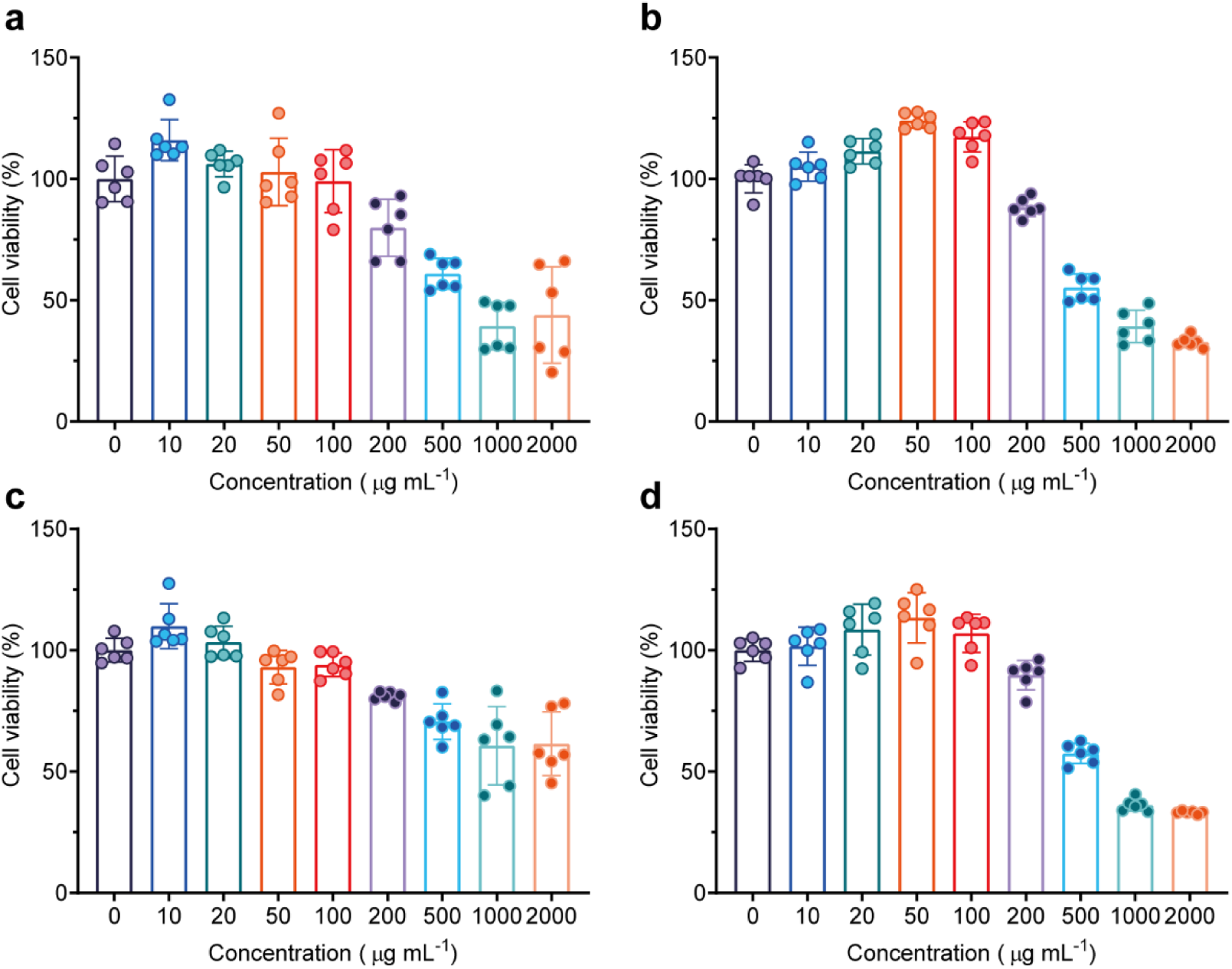
Cytotoxicity determined by CCK-8 assays. **a, b,** Viability of RAW264.7 cells (**a**) and NCM460 cells (**b**) after incubation with OPNs for 24 h. **c, d,** Viability of RAW264.7 cells (**c**) and NCM460 cells (**d**) after incubation with OPNs@LMWH for 24 h. Quantitative analysis of n=6 biologically independent samples (**a-d**) is shown. The data are presented as the means ± SDs.

**Supplementary Fig. 13.**
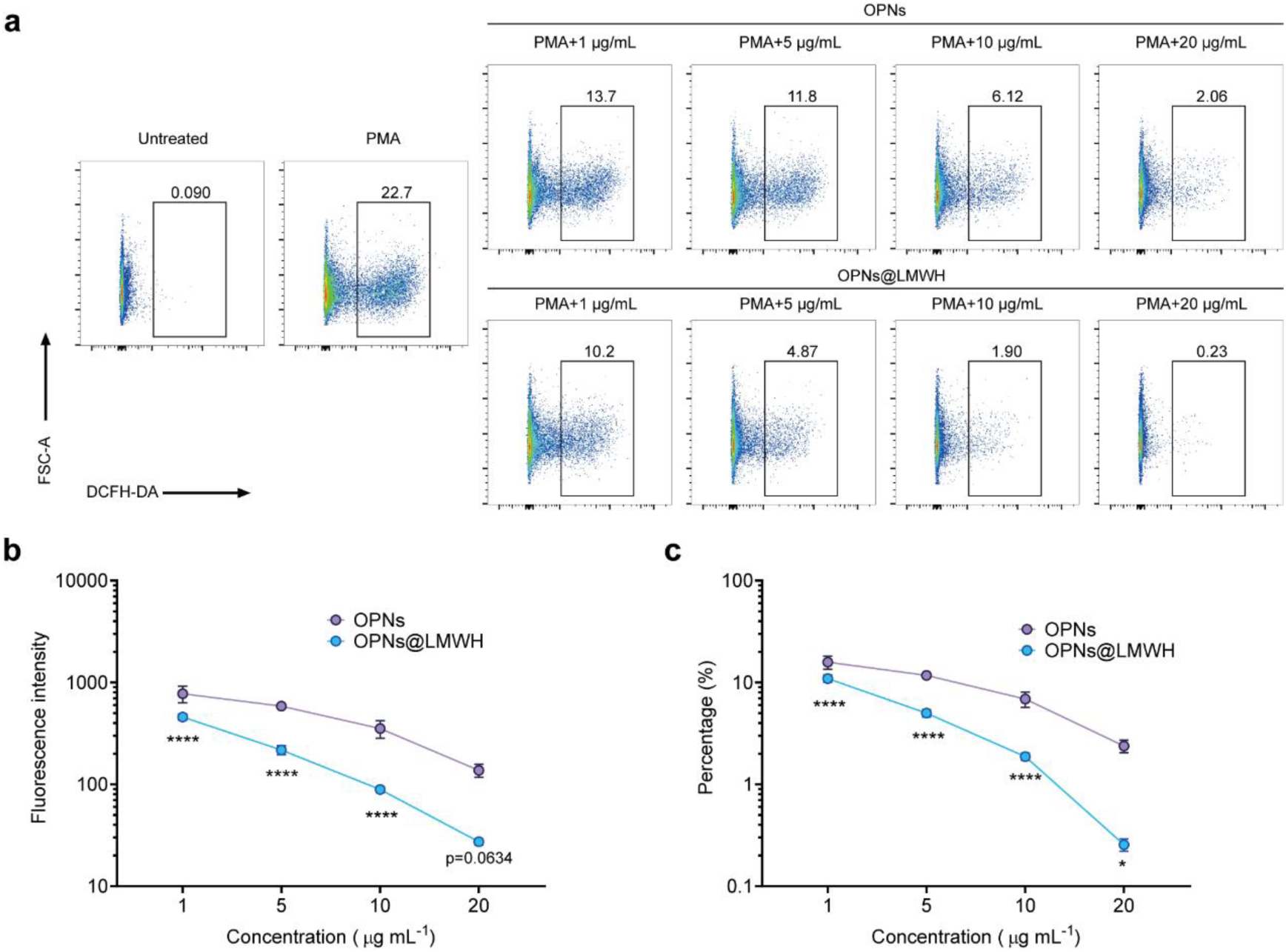
Concentration-dependent intracellular ROS scavenging behaviour. **a,** RAW264.7 cells were stimulated with PMA and treated with different concentrations of OPNs or OPNs@LMWH. **b, c,** Fluorescence intensities (**b**) and percentages of ROS-positive cells (**c**) were determined. The representative gating strategies or quantitative analyses of 4 biologically independent samples (**a-c**) are shown. The data are presented as the means ± SDs. * *p* < 0.05, ** *p* < 0.01, *** *p* < 0.001, **** *p* < 0.0001 and ns *p* > 0.05, analysed by two-way ANOVA (**b, c**).

**Supplementary Fig. 14.**
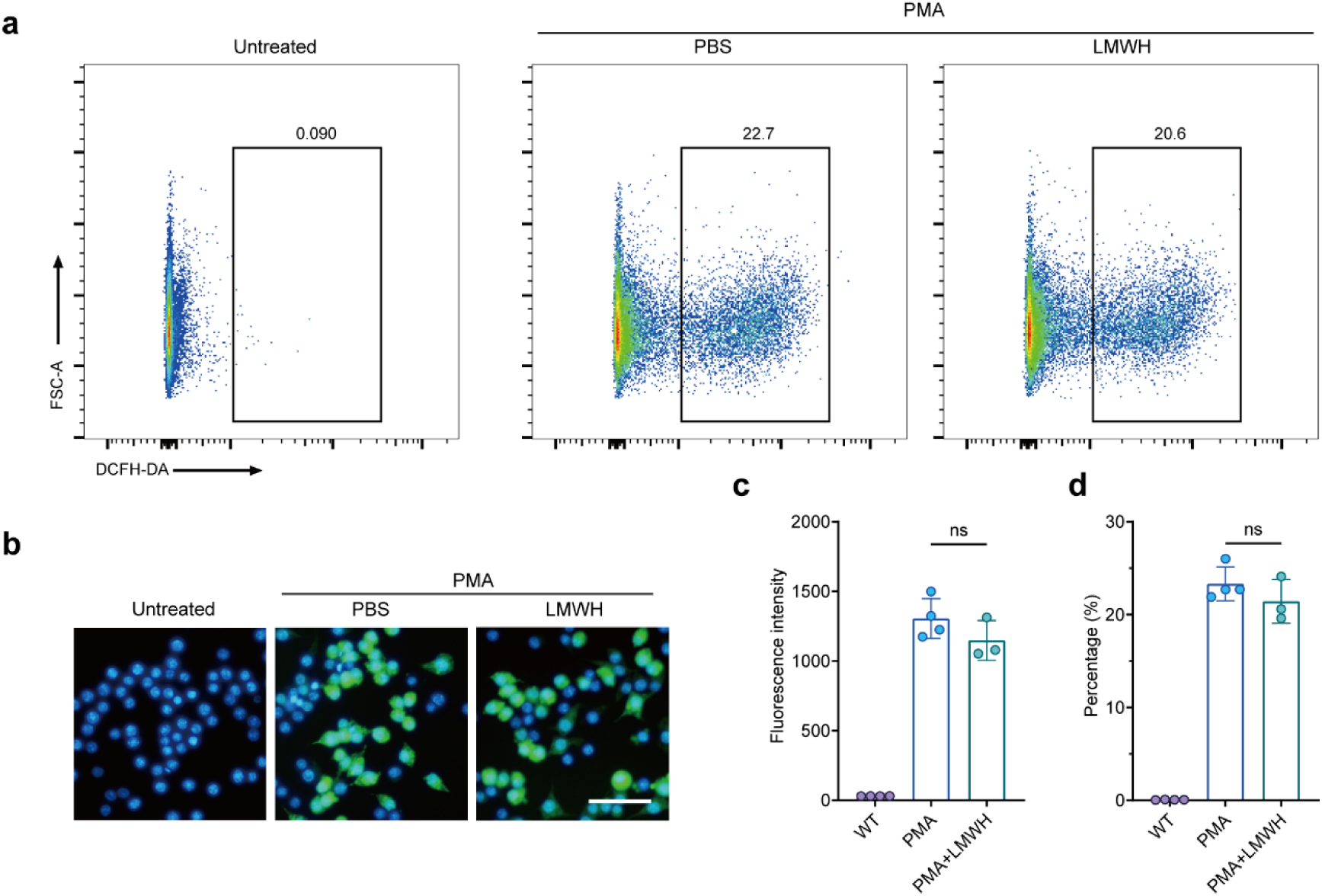
LMWH has little anti-ROS activity. **a, b,** RAW264.7 cells were stimulated with PMA and treated with LMWH (20 μg mL^-1^). The level of intracellular ROS was detected by DCFH-DA, determined by flow cytometry (**a**) and visualized by fluorescence microscopy (scale bar, 50 μm) (**b**). **c, d,** Fluorescence intensities (**c**) and percentages of ROS-positive cells (**d**) were determined. Representative gating strategies, images and quantitative analyses of n= 3 or 4 biologically independent samples (**a-d**) are shown. The data are presented as the means ± SDs. * *p* < 0.05, ** *p* < 0.01, *** *p* < 0.001, **** *p* < 0.0001 and ns *p* > 0.05, analysed by an unpaired t test (**c, d**).

**Supplementary Fig. 15.**
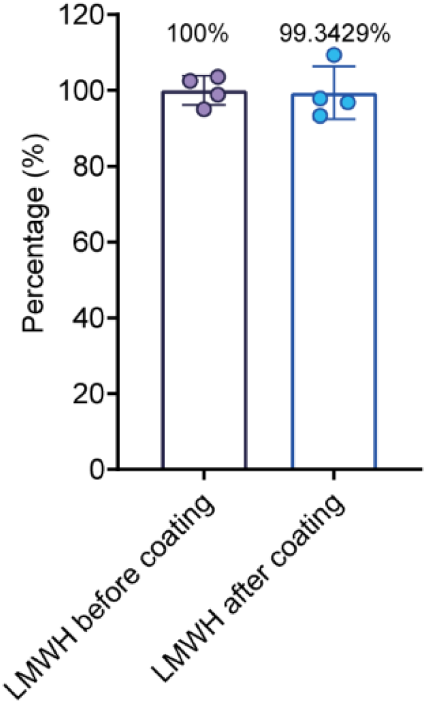
Concentration of free LMWH determined by Azure A. The amount of free LMWH in the supernatant before and after coating was detected, which indicated that LMWH was present in a low mass proportion in OPNs@LMWH. The quantitative analysis of 4 biologically independent samples is shown.

**Supplementary Fig. 16.**
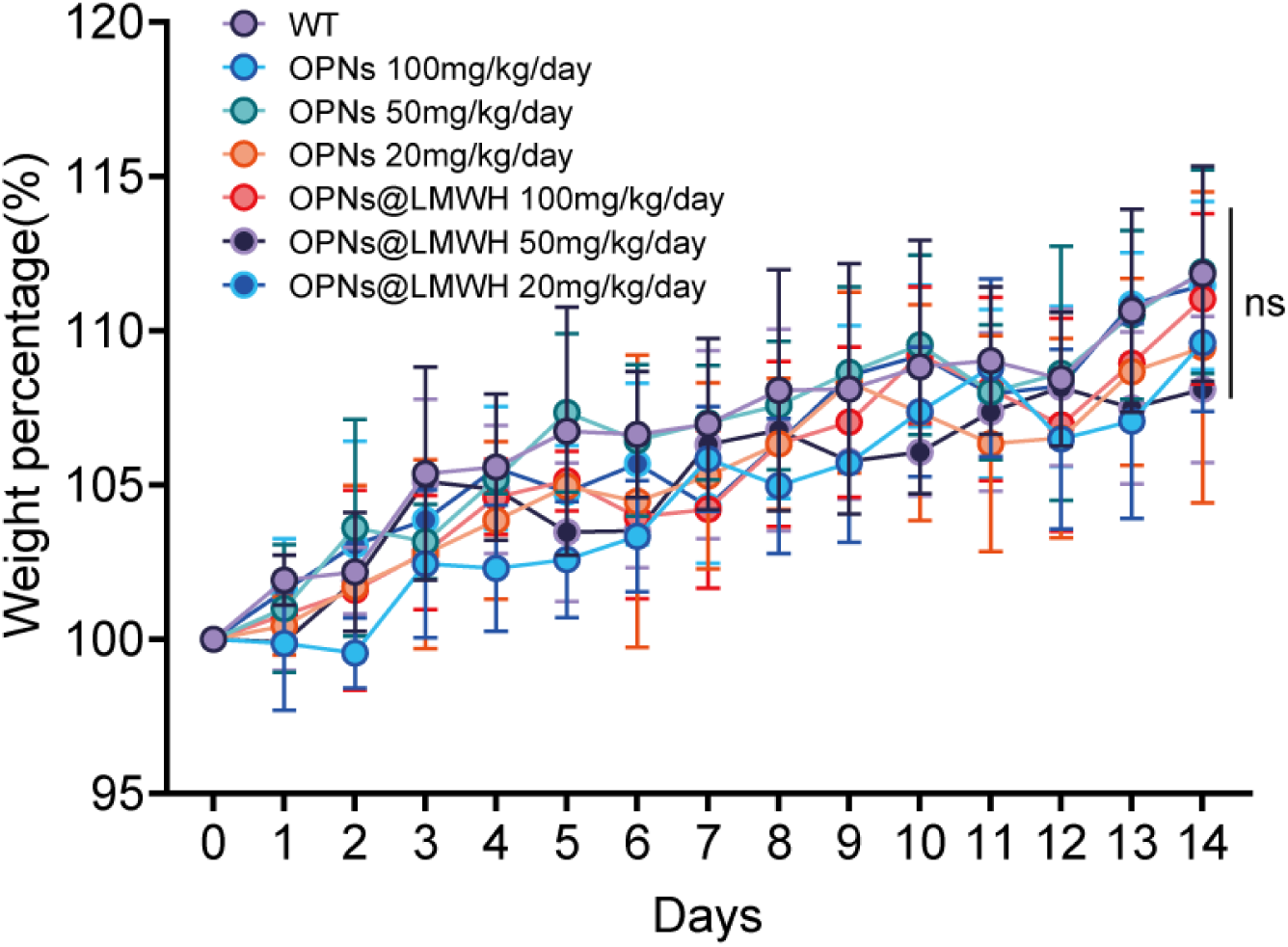
Safety profile of OPNs@LMWH (body weight). Mice were orally administered different doses of nanoparticles for 14 days. Daily body weight changes were recorded. Quantitative analysis of n=6 animals is shown. The data are presented as the means ± SDs. * *p* < 0.05, ** *p* < 0.01, *** *p* < 0.001, **** *p* < 0.0001 and ns *p* > 0.05, analysed by two-way ANOVA (**b, c**).

**Supplementary Fig. 17.**
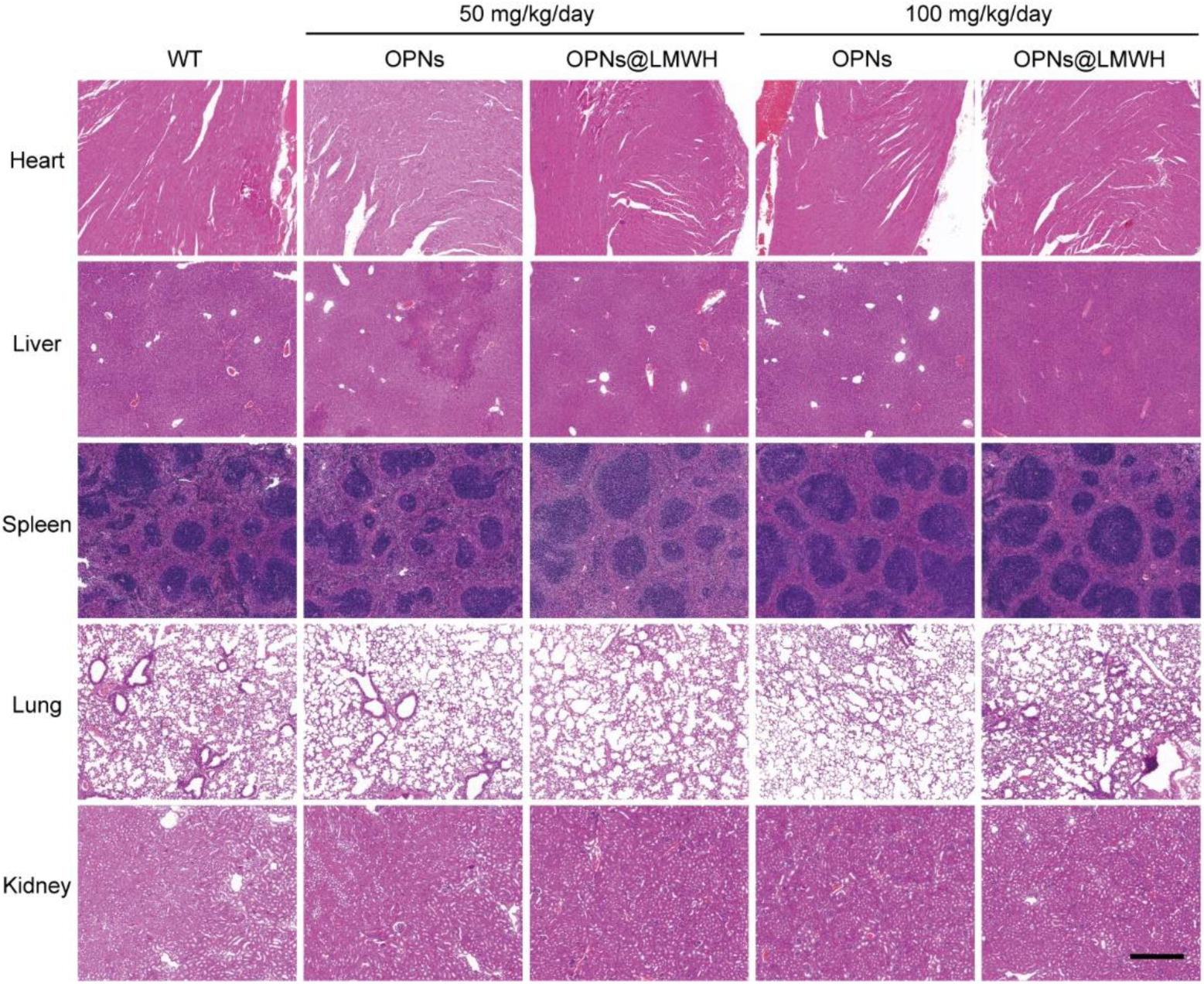
Safety profile of OPNs@LMWH (major organs). Mice were sacrificed on day 14. Major organ (heart, liver, kidney, lung, spleen, and colon) sections were stained with haematoxylin and eosin. Scale bar, 500 μm. Representative images from n= 3 animals are shown.

**Supplementary Fig. 18.**
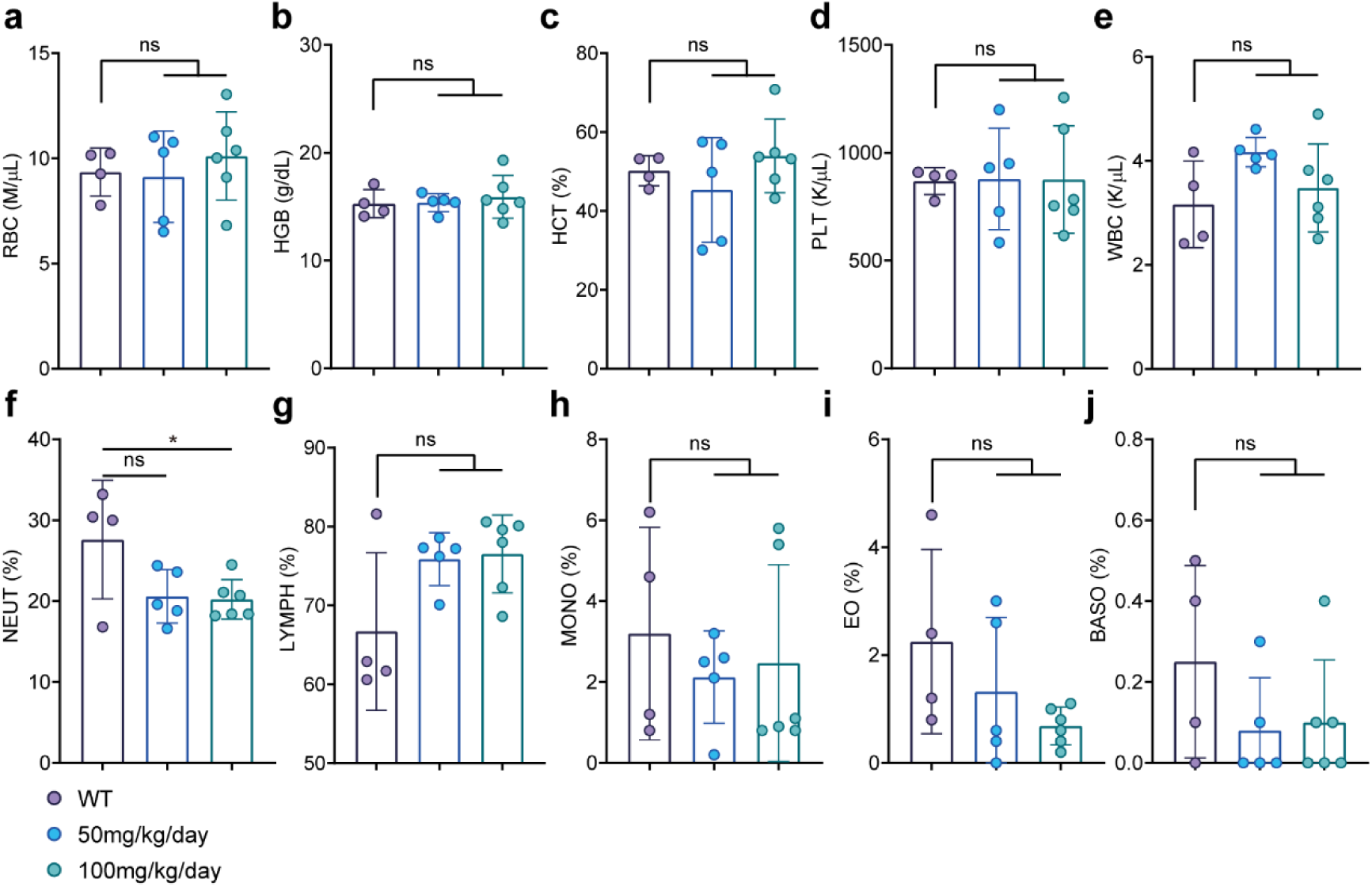
Safety profile of OPNs@LMWH (blood haematology). Mice were sacrificed on day 14. Blood samples were collected and analysed by using blood haematology; the results included red blood cell (RBC) (**a**), haemoglobin (HGB) (**b**), haematocrit (HCT) (**c**), platelet (PLT) (**d**), white blood cell (WBC) (**e**), neutrophil (NEUT) (**f**), lymphocyte (LYMPH) (**g**), monocyte (MONO) (**h**), eosinophil (EO) (**i**) and basophil (BASO) (**j**) counts. The quantitative analysis of n= 4-6 animals is shown. The data are presented as the means ± SDs. * *p* < 0.05, ** *p* < 0.01, *** *p* < 0.001, **** *p* < 0.0001 and ns *p* > 0.05, analysed by ordinary one-way ANOVA multiple comparisons test.

